# The COPI coatomer influences LDL receptor activity, hepatic lipid storage, and apoB secretion

**DOI:** 10.64898/2026.05.30.728950

**Authors:** Grigorios Panteloglou, Jérôme Robert, Marieke Smit, Nicolette Huijkman, Niels Kloosterhuis, Christopher S. Law, Brian Woods, Alaa Othman, Marcus E. Kleber, Graciela E. Delgado, Patrizia Tarugi, Museer A. Lone, Justina Clarinda Wolters, Antoine Rimbert, Anja Kerksiek, Dieter Lütjohann, Lucia Rohrer, Paolo Zanoni, Sofia Kakava, Stephanie Häusler, Eveline Schlumpf, Marta Futema, Steve E. Humphries, Janet Chou, Winfried März, Raif S. Geha, Anthony K. Shum, Jan Albert Kuivenhoven, Bart van de Sluis, Arnold von Eckardstein

## Abstract

**Background:** Decreased hepatic removal of low density lipoproteins (LDL) and increased apolipoprotein B (apoB) production cause hypercholesterolemia, a major causal risk factor of atherosclerotic cardiovascular disease (ASCVD). By a genome-wide siRNA screen, we previously identified subunits of the Coat protein I (COPI) complex to limit LDL uptake into Huh-7 hepatocarcinoma cells.

**Methods:** These findings were validated by targeted *in vitro* experiments as well as genetic association studies in humans and three mouse models with mutated or disrupted COPI genes.

**Results:** Silencing of *COPA*, *COPB1*, *COPB2*, *ARCN1*, *COPG1*, and *COPZ1* in Huh-7 cells resulted in decreased uptake of LDL and aberrant glycosylation and altered cell surface abundance of the LDL receptor (LDLR) as well as increased apoB secretion and cellular lipid storage. Single nucleotide polymorphisms of *ARCN1* were associated with lower ARCN1 expression and higher levels of LDL-cholesterol (LDL-C). Rare variants of *COPA* and *COPG1* were enriched among patients with LDL-C > 5 mmol/L. Patients and mice carrying other rare immunopathogenic missense variants of *COPA* and *COPG1* did not present with elevated plasma levels of LDL-C, while hepatic knockdown of murine *Copg1* increased the concentrations of non-HDL-cholesterol in plasma and triglycerides in the liver.

**Conclusions:** The COPI coatomer regulates LDLR activity and apoB secretion as well as lipid content of liver cells. Loss of function of some variants of COPI genes are associated with higher LDL-C levels.

## Introduction

The causal relationship between plasma levels of LDL cholesterol (LDL-C) and the risk of ASCVD is well-established^1^. Major determinants of LDL-C levels are the production of LDL’s canonical protein component apoB and the removal of LDL from the circulation by the liver through the binding of apoB to the LDL receptor (LDLR) for subsequent endocytosis and lysosomal degradation^2–4^. LDLR is a type 1 transmembrane protein comprised of 860 amino acid residues and synthesized in the endoplasmic reticulum (ER) as a precursor protein with an apparent molecular weight of 120 kDa^5^. After its production, LDLR is transported to the Golgi where both O- and N-glycosylation give rise to the mature LDLR isoform, which has an apparent molecular weight of 160 kDa^6^. The LDLR pathway is under tight regulation at multiple levels^2–4^. Its gene expression is induced by the sterol regulatory element binding protein 2 (SREBP2) and modulated by several microRNAs on the posttranscriptional level^2–4^. The U2-spliceosome enables the correct processing of transcripts that encode full length, functional LDLR^2–4^. In the presence of excess intracellular cholesterol, the Inducible Degrader of LDLR (IDOL) targets newly synthesized LDLR to lysosomal degradation. Ring finger containing protein 130 (RNF130) was recently identified as another E3 ubiquitin ligase that regulates hepatic LDLR at the post-translational level^3,4^. If taken up together with LDL, Proprotein Convertase Subtilisin Kexin 9 (PCSK9) directs LDLR-bound LDL to lysosomal degradation^3^. Otherwise LDLR is recycled to the cell surface by endosomes with the help of the Wiskott–Aldrich syndrome protein and SCAR homologue (WASH) and COMMD/CCDC22/CCDC93 (CCC) complexes^7,8^. Regulated Intramembrane Proteolysis (RIP) by metalloproteinase-mediated ectodomain shedding and subsequent γ-secretase-mediated cleavage of the membrane-bound stub of the receptor, also appears to regulate LDLR activity^3,4^. The extensive research of the LDLR pathway has led to the development of several LDL-C lowering drugs including statins, ezetimibe, bempedoic acid, and PCSK9 inhibitors, which also reduce the risk of cardiovascular events^1^.

Loss Of Function (LOF) variants in *LDLR* or, less commonly, specific missense variants in *APOB,* as well as very rare Gain-of-Function (GOF) variants in *PCSK9* cause autosomal dominant familial hypercholesterolemia (FH)^9^. Heterozygous FH (HeFH) is one of the most prevalent monogenic diseases with an estimated frequency of around 1/250 in many populations and, if untreated, leads to premature manifestation of ASCVD^10^. However, ∼40% of individuals with clinically diagnosed FH do not have any causal mutation in these genes, suggesting the involvement of still unknown genes in the regulation of the LDLR pathway^9^. To identify novel genes limiting hepatic LDL endocytosis, we previously performed a genome-wide siRNA screen in Huh-7 hepatocarcinoma cells^11^. The genes that significantly reduced LDL uptake upon knockdown included *COPA*, *COPB1,* and *COPB2,* which encode for three out of the seven core components of the COPI coatomer complex, namely α-COP, β-COP and β’-COP, respectively^11^. In another genome-wide RNAi screen conducted by Kraehling and colleagues, the knockdown of *ARCN1* encoding the δ-COP subunit of the COPI coatomer, also inhibited the uptake of LDL into the endothelial cell line EAhy.926^12^. The COPI coatomer coats vesicles mediating the retrograde and anterograde transport of proteins from the Golgi apparatus to the endoplasmic reticulum and from the *cis*- to the *trans*-Golgi network, respectively. It thereby plays an important role in the processing and quality control of proteins^13^. In addition, COPI contributes to the maturation of early endosomes, to lysosomal trafficking and autophagy^14,15^. Rare variants in *COPA, COPG1,* and *COPZ1* underlie inherited diseases characterized by autoinflammation, immunodeficiency, and neutropenia, respectively^16–19^ while certain missense mutations in *ARCN1* disturb craniofacial development^20^ . Because little is known about the impact of the COPI coatomer in lipoprotein metabolism. we decided to validate the findings of our siRNA screen by investigating the cellular mechanisms underlying disturbed lipoprotein uptake upon attenuated COPI function as well as the impact of genetic variants in COPI genes on plasma lipoprotein concentrations in humans and mice.

## Methods

### Data Availability

The authors declare that all data and methods supporting the findings of this study are reported in the main manuscript and in the Supplemental Material or are available from the corresponding authors on reasonable request.

### Human genetic studies

The associations of COPI gene polymorphisms and exome variants with lipid phenotypes were analysed by using the data banks of the Global Lipids Genetics Consortium^21^ and UK-Biobank ((https://www.azphewas.com: Dataset: UK Biobank 500k WGS (v2) Public)^22^ as reported previously^11,23^. The association of *ARCN1* rs17122278 alleles with the mRNA expression of *ARCN1* was analysed by using the databank of the Genotype-Tissue Expression (GTEx) project^24^.

Five patients with the COPA syndrome due to heterozygosity for the rare mutations in the *COPA* gene presented in figure 3C as well as the patients with combined immunodeficiency due to homozygosity for the rare p.K652E mutation in the *COPG1* gene and there heterozygous parents presented in figure 3D as well as the reference populations for comparison of LDLC were previously described^17,18,23,25–27^. Other rare *COPA* and *COPG1* variants presented in table 1 were identified by whole exome sequencing (WES) of patients with familial hypercholesterolemia (FH) who had no causative variant in the major FH-causing genes *LDLR*, *APOB*, and *PCSK9:*

**Table 1:**
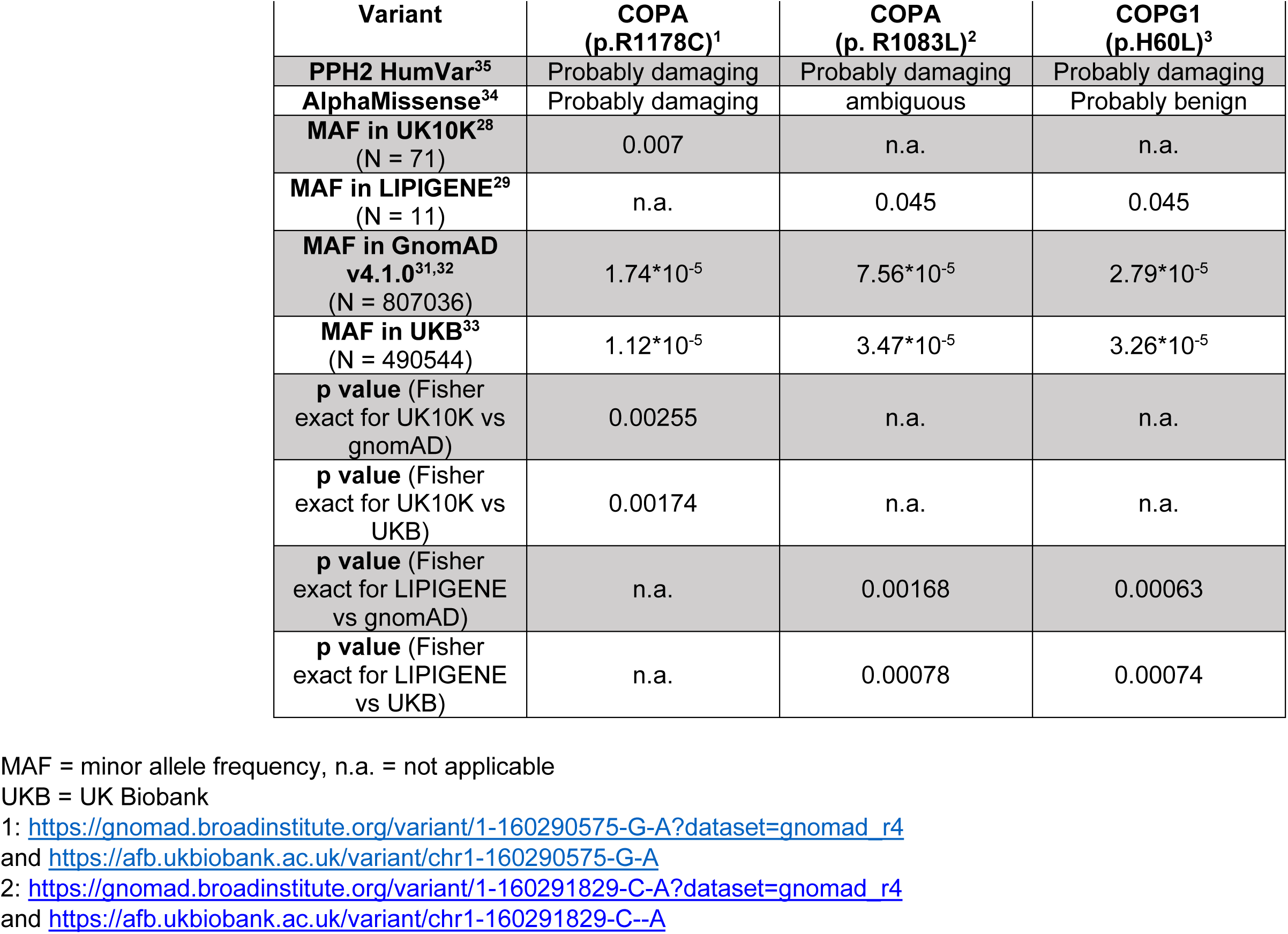
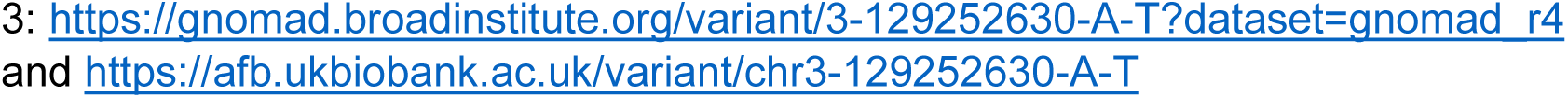
Variants identified in COPI genes of patients with familial hypercholesterolemia (FH) who have no causal variant in the canonical FH genes.

| **Variant** | **COPA**<br>**(p.R1178C)<sup>1</sup>** | **COPA**<br>**(p. R1083L)<sup>2</sup>** | **COPG1**<br>**(p.H60L)<sup>3</sup>** |
| --- | --- | --- | --- |
| **PPH2 HumVar<sup>35</sup>** | Probably damaging | Probably damaging | Probably damaging |
| **AlphaMissense<sup>34</sup>** | Probably damaging | ambiguous | Probably benign |
| **MAF in UK10K<sup>28</sup>**<br>(N = 71) | 0.007 | n.a. | n.a. |
| **MAF in LIPIGENE<sup>29</sup>**<br>(N = 11) | n.a. | 0.045 | 0.045 |
| **MAF in GnomAD**<br>**v4.1.0<sup>31,32</sup>**<br>(N = 807036) | $1.74 \times 10^{-5}$ | $7.56 \times 10^{-5}$ | $2.79 \times 10^{-5}$ |
| **MAF in UKB<sup>33</sup>**<br>(N = 490544) | $1.12 \times 10^{-5}$ | $3.47 \times 10^{-5}$ | $3.26 \times 10^{-5}$ |
| **p value** (Fisher<br>exact for UK10K vs<br>gnomAD) | 0.00255 | n.a. | n.a. |
| **p value** (Fisher<br>exact for UK10K vs<br>UKB) | 0.00174 | n.a. | n.a. |
| **p value** (Fisher<br>exact for LIPIGENE<br>vs gnomAD) | n.a. | 0.00168 | 0.00063 |
| **p value** (Fisher<br>exact for LIPIGENE<br>vs UKB) | n.a. | 0.00078 | 0.00074 |

MAF = minor allele frequency, n.a. = not applicable
UKB = UK Biobank
1: https://gnomad.broadinstitute.org/variant/1-160290575-G-A?dataset=gnomad_r4
and https://afb.ukbiobank.ac.uk/variant/chr1-160290575-G-A
2: https://gnomad.broadinstitute.org/variant/1-160291829-C-A?dataset=gnomad_r4
and https://afb.ukbiobank.ac.uk/variant/chr1-160291829-C--A
3: https://gnomad.broadinstitute.org/variant/3-129252630-A-T?dataset=gnomad_r4 and https://afb.ukbiobank.ac.uk/variant/chr3-129252630-A-T

COPA(p.R1178C) (GRCh38: chr1:160290575G>A) variant was found as part of the UK10K project^28^ while the COPA p.R1083L (GRCh38: chr1-160291829-C-A) and COPG1 p.H60L/chr3-129252630-A-T variants were found in the UNIMORE-FH LIPIGEN project^29^. Six other rare variants of *COPA* or *COPG1* presented in table 2 were identified by WES using DNA from 96 participants in the LURIC study^30^ who had LDL-cholesterol levels >190 mg/dL either actually or likely before treatment with lipid lowering drugs. We compared the minor allele frequencies of these variants with those reported in the databanks of GnomADv4.1.0^31,32^ (n=807,036) and UK Biobank (n=490,544)^33^. For the prediction of pathogenicity we used AlphaMissense^34^ and PolyPhen2^35^.

**Table 2.**
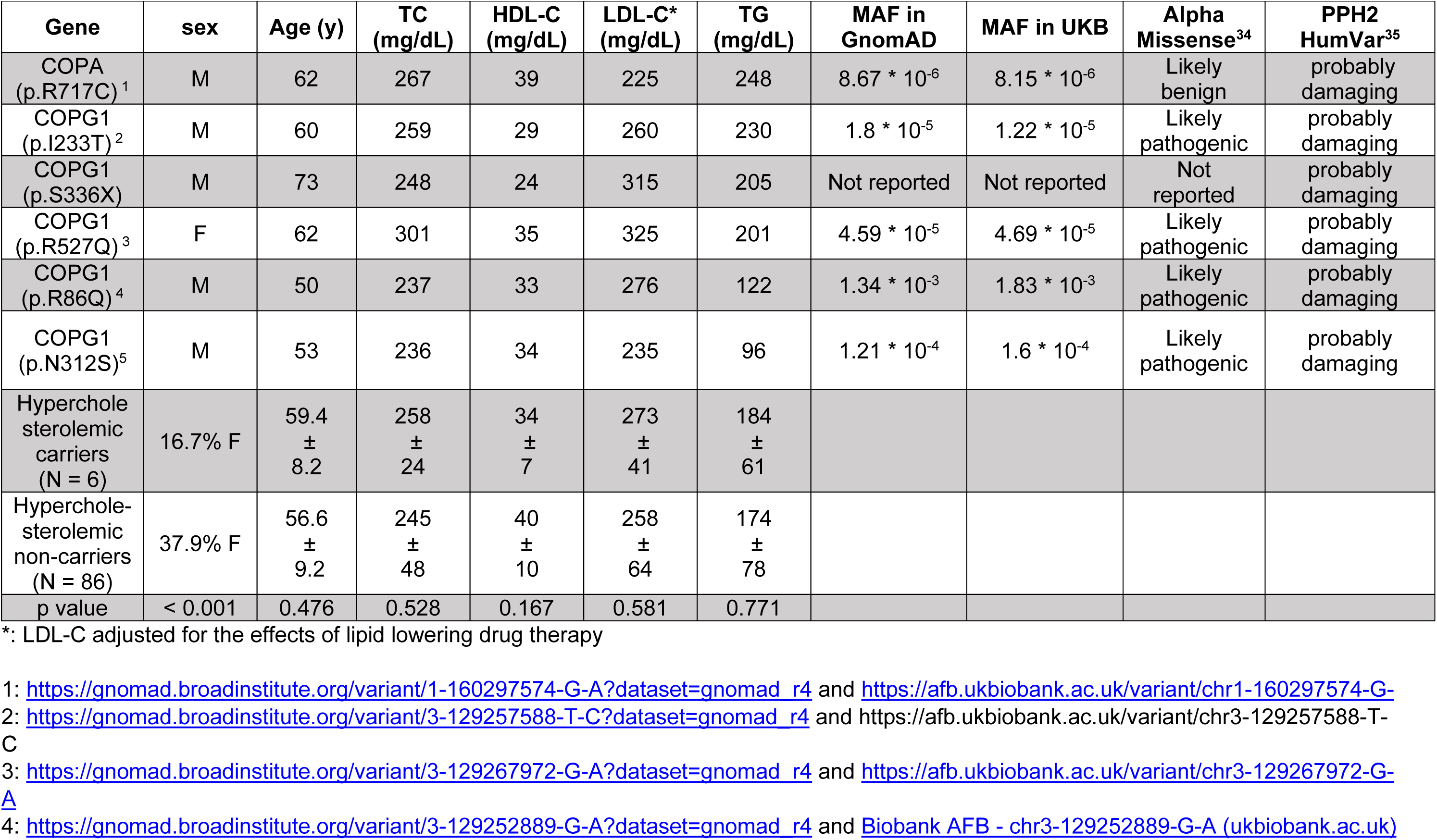
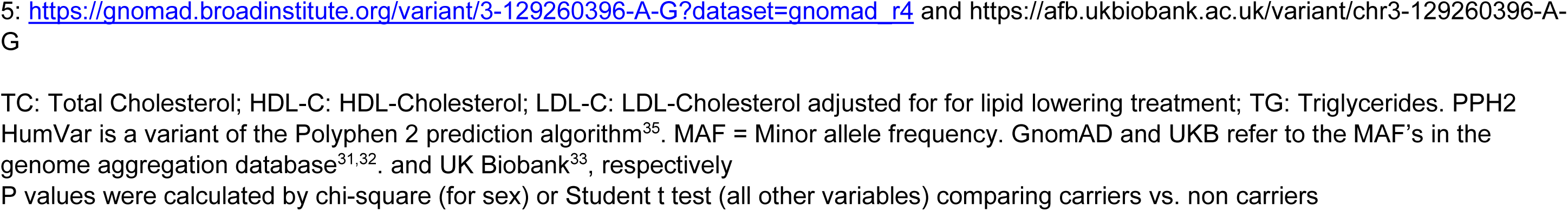
Rare variants identified in COPI genes in hypercholesterolemic participants of the LURIC ^30^ study.

| Gene | sex | Age (y) | TC (mg/dL) | HDL-C (mg/dL) | LDL-C* (mg/dL) | TG (mg/dL) | MAF in GnomAD | MAF in UKB | Alpha Missense<sup>34</sup> | PPH2 HumVar<sup>35</sup> |
| --- | --- | --- | --- | --- | --- | --- | --- | --- | --- | --- |
| COPA (p.R717C)<sup>1</sup> | M | 62 | 267 | 39 | 225 | 248 | 8.67 * 10<sup>-6</sup> | 8.15 * 10<sup>-6</sup> | Likely benign | probably damaging |
| COPG1 (p.I233T)<sup>2</sup> | M | 60 | 259 | 29 | 260 | 230 | 1.8 * 10<sup>-5</sup> | 1.22 * 10<sup>-5</sup> | Likely pathogenic | probably damaging |
| COPG1 (p.S336X) | M | 73 | 248 | 24 | 315 | 205 | Not reported | Not reported | Not reported | probably damaging |
| COPG1 (p.R527Q)<sup>3</sup> | F | 62 | 301 | 35 | 325 | 201 | 4.59 * 10<sup>-5</sup> | 4.69 * 10<sup>-5</sup> | Likely pathogenic | probably damaging |
| COPG1 (p.R86Q)<sup>4</sup> | M | 50 | 237 | 33 | 276 | 122 | 1.34 * 10<sup>-3</sup> | 1.83 * 10<sup>-3</sup> | Likely pathogenic | probably damaging |
| COPG1 (p.N312S)<sup>5</sup> | M | 53 | 236 | 34 | 235 | 96 | 1.21 * 10<sup>-4</sup> | 1.6 * 10<sup>-4</sup> | Likely pathogenic | probably damaging |
| Hypercholesterolemic carriers (N = 6) | 16.7% F | 59.4 ± 8.2 | 258 ± 24 | 34 ± 7 | 273 ± 41 | 184 ± 61 |  |  |  |  |
| Hypercholesterolemic non-carriers (N = 86) | 37.9% F | 56.6 ± 9.2 | 245 ± 48 | 40 ± 10 | 258 ± 64 | 174 ± 78 |  |  |  |  |
| p value | < 0.001 | 0.476 | 0.528 | 0.167 | 0.581 | 0.771 |  |  |  |  |

\*: LDL-C adjusted for the effects of lipid lowering drug therapy
- 1: https://gnomad.broadinstitute.org/variant/1-160297574-G-A?dataset=gnomad_r4 and https://afb.ukbiobank.ac.uk/variant/chr1-160297574-G-A - 2: https://gnomad.broadinstitute.org/variant/3-129257588-T-C?dataset=gnomad_r4 and https://afb.ukbiobank.ac.uk/variant/chr3-129257588-T-C - 3: https://gnomad.broadinstitute.org/variant/3-129267972-G-A?dataset=gnomad_r4 and https://afb.ukbiobank.ac.uk/variant/chr3-129267972-G-A - 4: https://gnomad.broadinstitute.org/variant/3-129252889-G-A?dataset=gnomad_r4 and [Biobank AFB - chr3-129252889-G-A \(ukbiobank.ac.uk\)](https://afb.ukbiobank.ac.uk/variant/chr3-129252889-G-A)
5: https://gnomad.broadinstitute.org/variant/3-129260396-A-G?dataset=gnomad_r4 and https://afb.ukbiobank.ac.uk/variant/chr3-129260396-A-G
TC: Total Cholesterol; HDL-C: HDL-Cholesterol; LDL-C: LDL-Cholesterol adjusted for lipid lowering treatment; TG: Triglycerides. PPH2 HumVar is a variant of the Polyphen 2 prediction algorithm<sup>35</sup>. MAF = Minor allele frequency. GnomAD and UKB refer to the MAF's in the genome aggregation database<sup>31,32</sup>. and UK Biobank<sup>33</sup>, respectively
P values were calculated by chi-square (for sex) or Student t test (all other variables) comparing carriers vs. non carriers

### Animal studies

Heterozygous Copa^wt/E241K^ and homozygous Copg1^K652E/K652E^ mice have been previously described^17,23,36^. To generate mice with hepatocyte-specific Cas9 expression, Rosa26-LSL-Cas9 knck-in mice cat. 024857, The Jackson Laboratory)^37^ were crossed with Albumin-Cre mice (cat. 003574, The Jackson Laboratory). *Copg1* was targeted in mouse hepatocytes, using CRISPR/Cas9 gene editing technology^8^ using adeno-associated virus (AAV)^38^. Gene blocks, consisting of three single guide RNAs (sgRNAs) targeting *Copg1* were ordered (ThermoFisher Scientific); the complete sequences are available upon request. Male mice (littermates, 8-12 weeks of age at the time of gene editing) were used for all experiments and were individually housed under a 12-hour light-dark cycle (lights on at 8:00) with ad libitum access to standard laboratory diet (cat. V1554, Ssniff Spezialdiäten GmbH, Germany) and water. Prior to sacrifice, mice were fasted for 4 hours. Upon sacrifice, liver tissue was collected and snap-frozen in liquid nitrogen and stored at -80°C until further analysis. Blood was collected by cardiac puncture into EDTA-coated tubes, and plasma was collected after centrifugation at 1000 x g for 10 min at 4°C. Blood samples were collected into tubes containing EDTA while the livers of the mice were harvested after the cardiac puncture. For several analyses plasmas and liver tissues were frozen and shipped on dry ice to Zurich.

Concentrations of cholesterol, triglycerides and HDL-cholesterol as well as the activity of alanine amino transferase (ALT) in mouse plasmas were measured using photometric assays from Roche diagnostics on the COBAS8000 autoanalyser (Rotkreuz, Switzerland). NonHDL-cholesterol was calculated as the difference between total and HDL-C. Plasma concentrations of apoA-I and apoB were determined with immunonephelometric assays and the Attelica Neph 630 from Siemens Healthineers. Protein concentrations of total apoB (apoB_100_+apoB_48_) and apoB_100_, were also quantified by targeted proteomic assays as previously described^8,23^.

Lipids in homogenates of mouse liver (15% w/v in PBS) were extracted as principally described^39^. First, 200 μL of liver homogenate was added to 600 μL of demi-water, mixed with 3 mL chloroform/methanol (1:2 v/v) in a glass tube, and incubated for 30 minutes. Then, 1.2 mL of H_2_O and 1 mL of chloroform were added, mixed, and samples were centrifuged at 500 x g for 10 minutes at room temperature. The organic layer was transferred to a new glass tube, and the solvent was evaporated with nitrogen at 50°C. The dried lipids were dissolved in 1 mL chloroform, from which 400 μL was again dried and dissolved in 500 μL Triton X-100 2% in chloroform, and evaporated with nitrogen at 50°C. Then, 500 μL demi-water was added, and samples were incubated for 15 minutes at 37°C before further analysis for cholesterol and triglyceride concentrations. Total cholesterol and triglyceride concentrations in liver extracts were quantified with colorimetric assays from Roche cat. 11489232 and cat. 1187771, respectively) and standard FS (DiaSys) and Precimat Glycerol standard (Roche), respectively, as calibrators.

### Cell culture

Huh-7 cells (cat. JCRB0403, JCRB Cell Bank, Ibaraki, Japan) were cultured in normal growth medium as described previously^23^.

### siRNA transfection and overexpression of GFP-LDLR in Huh-7 cells

Huh-7 were reverse transfected with either siRNAs against the indicated target genes (*COPA*, *COPB1*, *COPB2*, *ARCN1*, *COPG1*, *COPG2*, *COPE*, *COPZ1*, *COPZ2, LDLR*) or non-targeting siRNAs (please see Major Resources Table for details) as described previously^23^.

The plasmid containing the coding sequence of *LDLR* (NM_000527) carrying an N-terminal GFP tag was purchased from addgene^40^ (cat. 98184, Watertown, USA). The plasmid transfection and the selection of cells successfully taking up the plasmids were carried out as previously described^23^. The cells were transfected with either an empty plasmid (cat. EX-NEG-M02, GeneCopoeia) alone (negative control Empty Vector cells) or a combination of the empty plasmid together with the plasmid encoding for GFP-LDLR (cells designated overexpressing GFP-LDLR)

### Real-Time Quantitative Reverse Transcription PCR (qRT-PCR)

Huh-7 cells were harvested, their RNA was isolated, reverse transcribed and the derived cDNA was used for qRT-PCR as described previously^23^. Sections of mouse livers were homogenized using Tri reagent® (cat. T9424, Sigma-Aldrich,) and the extracted RNA was treated in the same way as the cells. For each experiment, 3 technical replicates were used for each condition. At the end of each PCR run, the specificity of the PCR products was confirmed by melting temperature (Tm) analysis. The data derived from Huh-7 cells were analyzed by performing relative quantification based on crossing point (Cp) values for the reference gene (GAPDH) and the gene of interest using the relative standard curve method. For the data derived from mice for each condition, the ratio of the signal for each gene of interest / signal GAPDH was calculated and the data were normalized to the respective control condition, as described for each experiment. Primer sequences and their respective target genes are described in Supplemental Table S1.

### Uptake of fluorescently labeled LDL

LDL was isolated from human plasmas of normolipidemic donors by sequential ultracentrifugation and labeled with Atto655 (cat. AD655-35, Atto-Tec, Siegen, Germany) as reported previously^41^. 72 hours after transfection with the indicated siRNAs, the cells were incubated at 37°C with 10 μg/mL of Atto655-LDL in DMEM supplemented with 0.2% BSA in the presence (unspecific uptake) or absence (total uptake) of 100 times excess of unlabeled LDL. After 2 hours, the cells were extensively washed with PBS and detached by incubation with Accutase^®^ (cat. A6964, Sigma-Aldrich) for 5 minutes at 37°C. The samples were then processed and analysed by flow cytometry as described previously for the uptake of HDL using the BD LSR II Fortessa and BD FACSDIVA™ software (BD-Biosciences)^23^. Data analysis was carried out using FlowJo version 10 (FlowJo LLC, Ashland, USA). The Median Fluorescence Intensity (MFI) of each population was used for comparison between the different conditions and the specific uptake was calculated as the difference between total uptake and unspecific uptake for each condition.

### Subcellular localization of GFP-LDLR by flow cytometry or confocal microscopy

LDLR and GFP cell surface levels as well as the overall emitted GFP signal were determined by flow cytometry using alive wild-type (figures 1E and 1F) or Huh-7 cells overexpressing either an empty vector alone (EV) or in combination with a vector encoding for GFP-LDLR (GFP-LDLR) (supplemental figures 3C and 3D display experiments with both EV and GFP-LDLR, while figures 2A-2C experiments with only cells overexpressing GFP-LDLR). 72 hours after transfection or 56 hours after transfection and 16 hours after treatment with DMSO (see legends of the figures for the specific condition), the medium was aspirated and the cells were detached and prepared for investigation of cell surface LDLR or GFP levels or overall GFP levels as described previously^23^. Deviations from the previously described protocol included the treatment of Huh-7 cells overexpressing the empty vector alone or in combination with the plasmid encoding for GFP-LDLR with PI only for the exclusion of signal originating from dead cells, and the lack of compensation for spectral overlap in experiments measuring GFP cell surface levels, due to the inability of having single-stained cells for the fluorophore assigned to the respective secondary antibody without having GFP signal present. Sources and concentrations of both primary and secondary antibodies used are summarized in the Major Resources Table.

**Figure 1.**
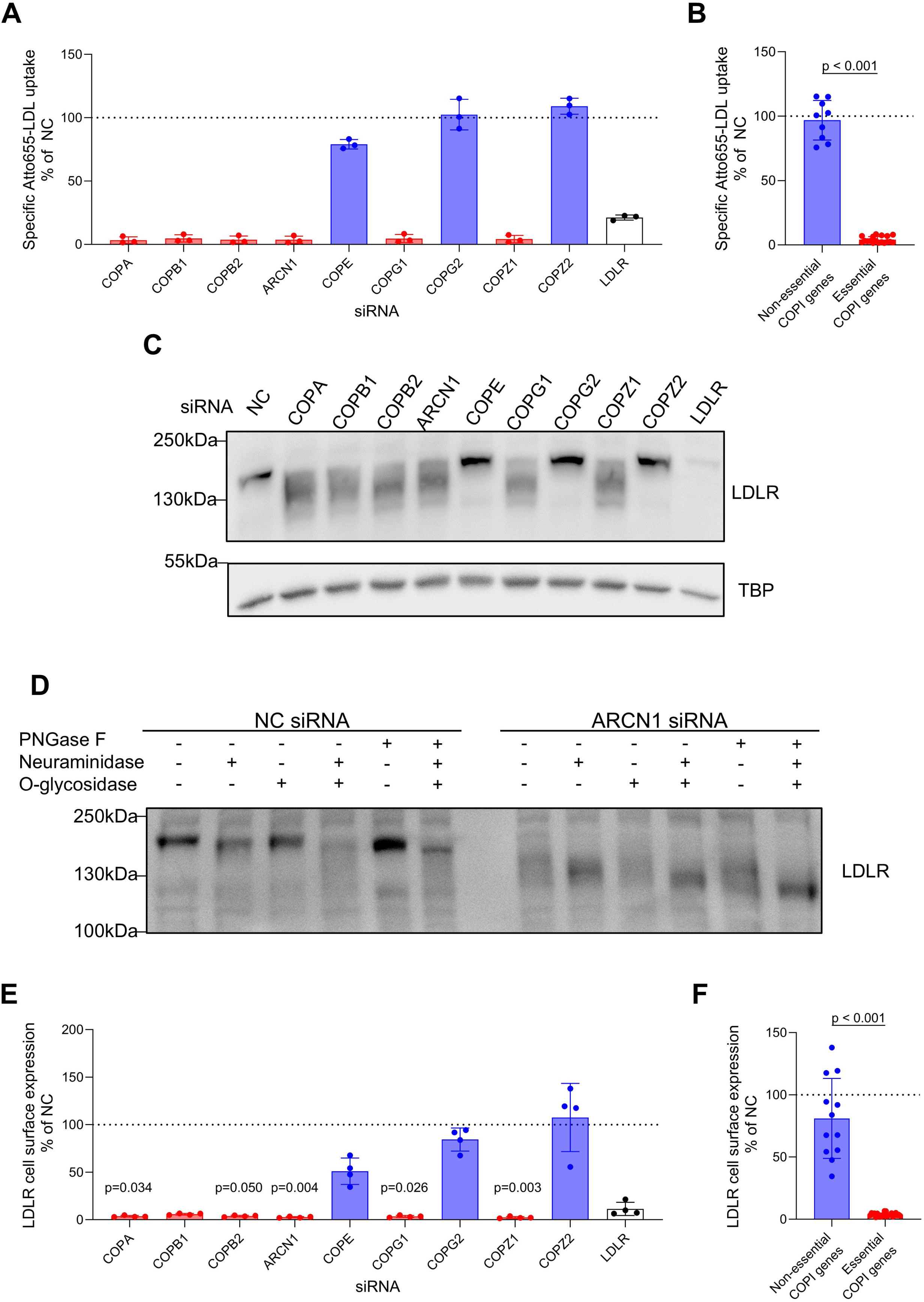
Silencing of the COPI subunits reduces LDL uptake and alters the electrophoretic properties and the cell surface abundance of LDLR in Huh-7 hepatocarcinoma cells. Wild type Huh-7 cells were transfected with the indicated siRNAs (**A**-**F**) and 72 hours post transfection were used for functional evaluation. **A** and **B**. The cells were incubated with 10 μg/mL Atto655-LDL for 2 hours in the presence or absence of 100-fold excess of unlabeled LDL and then collected for measurement with flow cytometry. The specific cell association was calculated as the difference between the two conditions. The data were normalized to the non-coding control (NC). **C** and **D** show representative Western Blots out of 3 (**C**) or 2 (**D**) independent experiments using lysates immediately after harvesting (**C**) or after overnight treatment with the indicated combinations of deglycosylating enzymes (**D**). LDLR was immunodetected with an antibody against an intracellular, C-terminal epitope and TATA binding protein (TBP) served as a loading control in (**C**). Figures **E** and **F** show LDLR cell surface levels of alive Huh-7 cells as determined with an antibody against an extracellular, N-terminal epitope of LDLR. All data (**A**, **B**, **E**, **F**) were normalized to the non-coding (NC) control condition and are shown as means ± SD of three (**A**, **B**) or four (**E**, **F**) independent experiments. **B** and **F** compare merged data on the indispensable COPI subunits (red bars) and the dispensable or paralogous COPI genes (blue bars). Statistical analysis was performed using Kruskal-Wallis test with Dunn’s multiple comparisons test between NC and each targeting siRNA (**A**, **E**) or Mann-Whitney test (two-tailed) (**B**) or unpaired t-test (two-tailed) (**F**) between the essential and the non-essential conditions. Only statistically significant differences (p<0.05) are indicated.

**Figure 2.**
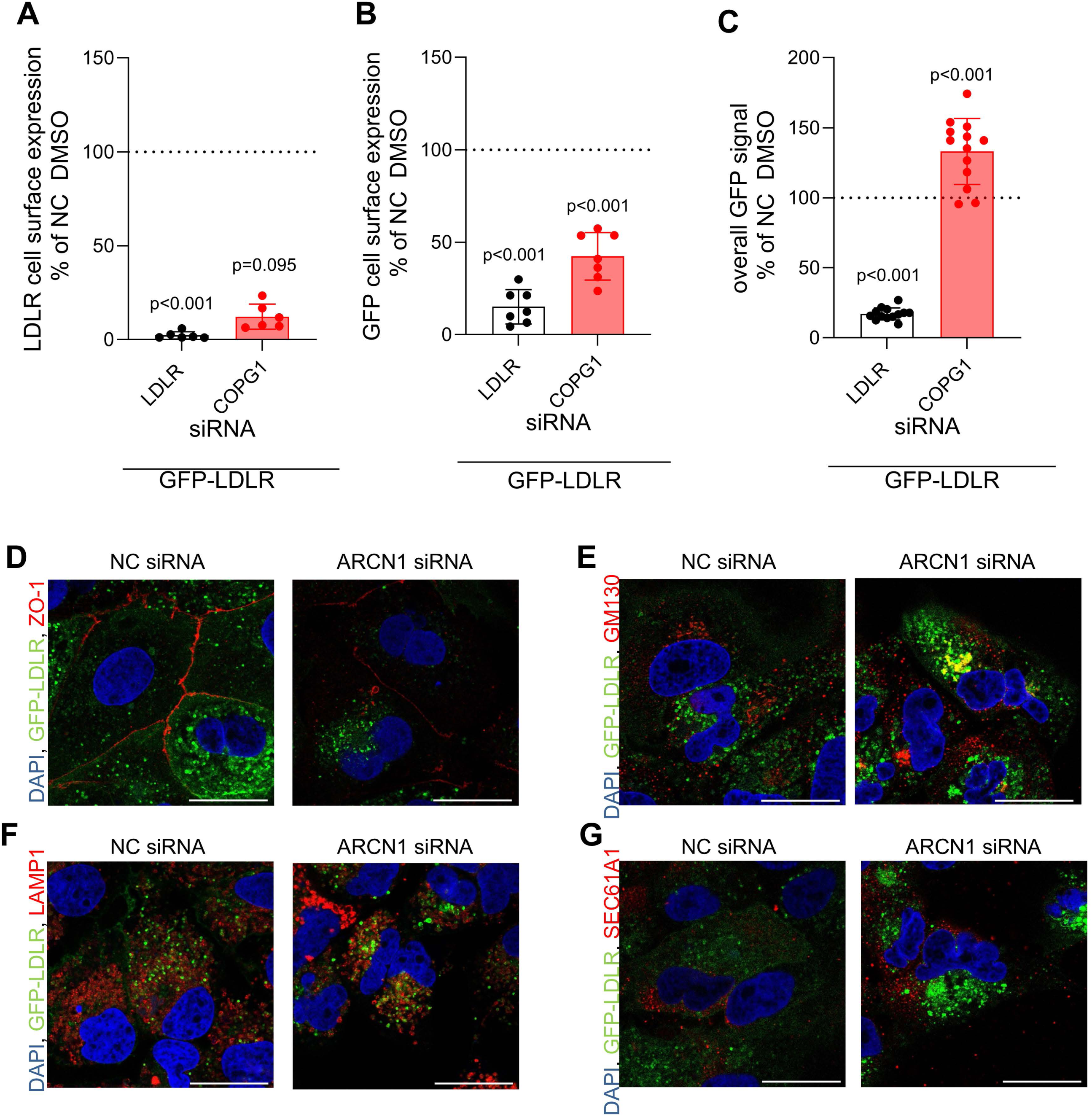
Silencing of COPI subunits leads to intracellular accumulation of LDLR. Huh-7 cells stably overexpressing an empty vector together with another vector containing *LDLR* with a GFP tag attached to its aminoterminal end (GFP-LDLR) were transfected with the indicated siRNAs (**A**-**G**). 56 hours after transfection followed by 16 hours treatment with DMSO (**A**-**C**), or 72 hours after transfection (**D-G**) the cells were collected and used for the analysis of LDLR. Figure parts (**A, B**) show the flow cytometry-based measurement of LDLR (**A**) or GFP (**B**) cell surface levels using either an antibody against an aminoterminal epitope of LDLR or an anti-GFP antibody, respectively. The fluorescence emitted by the GFP-LDLR was recorded as total cellular LDLR (**C**). The data were normalized to the NC DMSO condition and are shown as means ± SD of six (**A),** seven **(B**) or thirteen (**C**) independent experiments. Statistical analyses were performed using either Kruskal-Wallis test with Dunn’s test for multiple comparisons (**A**) or 1-way ANOVA coupled with Dunnett’s test for multiple comparisons (**B**, **C**) between NC and each targeting siRNA. Figure parts **D-G** depict confocal microphotographs on the colocalization of GFP-LDLR with ZO-1 (**D**), GM130 (**E**), LAMP1 (**F**) and SEC61A1 (**G**) in Huh-7 cells overexpressing GFP-LDLR, 72 hours after transfection with the indicated siRNAs. The data shown are representative of two independent experiments. The scale bar for both microphotographs is 25 μm.

**Figure 3.**
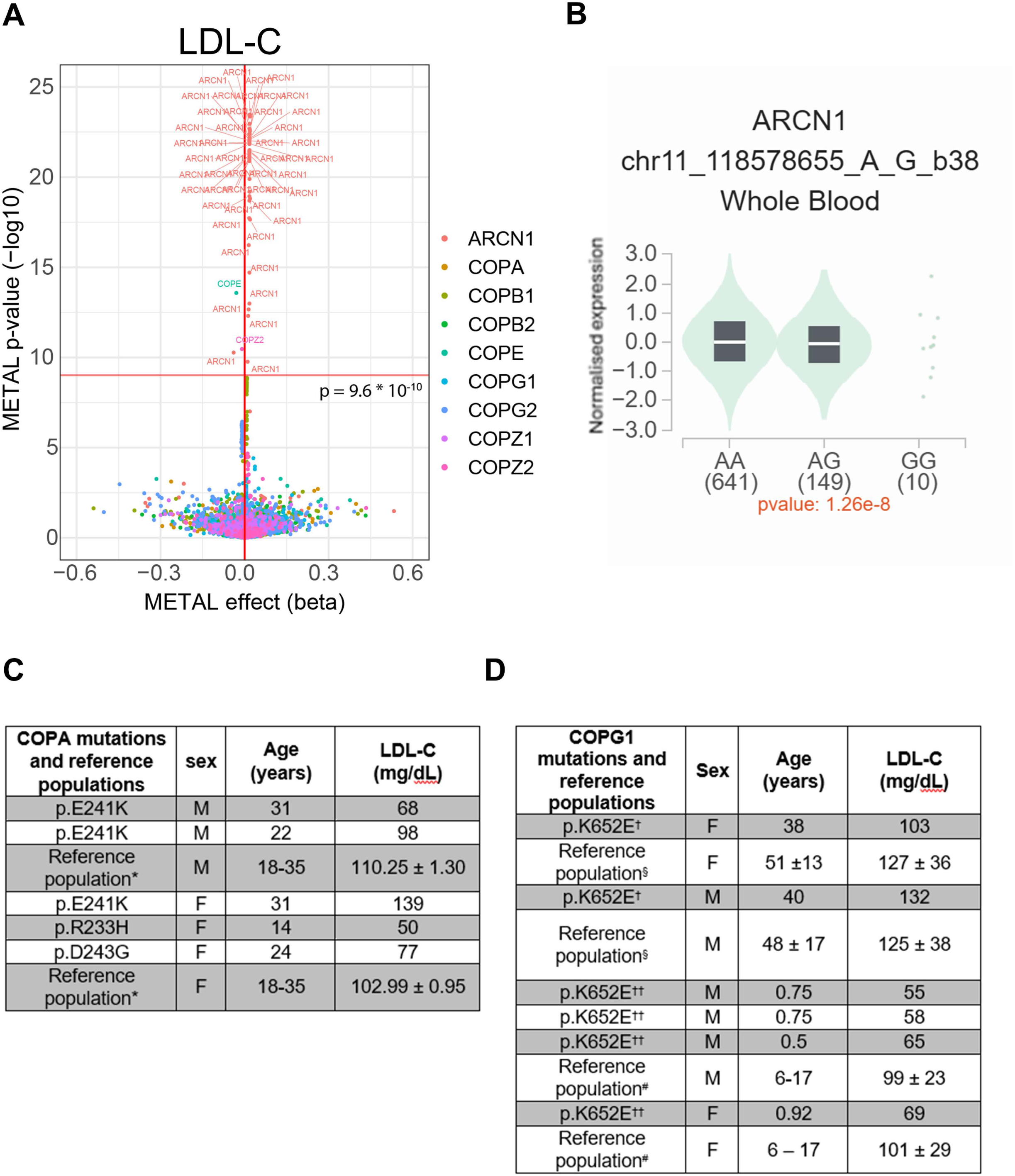
Associations of SNPs (A, B) or rare variants (C, D) of COPI genes with plasma levels of LDL-C (A, C, D) or ARCN1 mRNA expression by blood cells (B) in humans. **A.** Volcano plot depicting the associations of 5,569 SNPs of the nine COPI genes with LDL-C in 1.65 million individuals aggregated by the Global Lipid Genetics Consortium^21^. The x axis depicts the effect size (beta) and the y-axis the p-value. The dashed horizontal line marks the threshold of statistical significance after Bonferroni correction for multiple testing on GWAS level (p = 9.60*10^−10^). The violin plot (**B**) shows the association of the ARNC1 rs11216916 alleles with the mRNA expression of *ARCN1* in blood of nearly 1000 individuals collected by the Genotype Tissue Expression (GTEx) project^24^ (p=6.01 *10^−10^, NES = -0.12). **C** and **D** show LDL-C levels in individuals suffering from COPA syndrome^18^ and COPG1 syndrome^17^, respectively. For comparison with the US-American COPA variant carriers (**C**), data of 6,127 male and 6,157 female participants of NHANES^25^ are presented (*) as means± standard error. For comparison of the heterozygous (†) and homozygous (††) carriers of the rare COPG1 variant (**D**) who originate from Oman, data of adults (§) and children and adolescents (#) from the United Arab Emirates are shown for comparison^26,27^. Data are means ± SD of 485 men and 492 women (§) or 490 boys and 476 girls (#). LDL-C: LDL-Cholesterol; SD: Standard deviation.

GFP-LDLR was colocalized with organelle markers in Huh-7 cells overexpressing GFP-LDLR by the use of confocal microscopy principally as described previously for the colocalization of GFP-SR-BI^23^. Sources and concentrations of both primary and secondary antibodies used are summarized in the Major Resources Table.

### Western Blotting

Lysates of Huh-7 cells were obtained, processed and used for Western Blotting as described previously^23^. The intensity of the bands was recorded by densitometry using ImageJ^42^. Sources and concentrations of both primary and secondary antibodies used are summarized in the Major Resources Table.

### Enzymatic deglycosylation

Lysates of Huh-7 cells transfected with the indicated siRNAs were used for enzymatic deglycosylation as described previously^23^.

### Quantification of apoB secreted by Huh-7 cells

Newly synthesized and secreted apoB was measured by the incorporation of 35S-methionine and cysteine followed by immunoprecipitation. Briefly, Huh-7 cells were seeded into 6-well plates and transfected with siRNAs, as described above. After 72 hours, culture media was removed and cells were placed in DMEM without methionine and cysteine but with 1.5% of BSA. After 2 hours, cells were labeled with 150 μCi/mL of ^35^S methionine and cysteine (Hartman, Heidenheim Germany) in the same media. Antibodies against apoB were added to the media and incubated at 4°C with constant mixing for at least 2 hours. Protein A/G magnetic agarose beads (cat. 78609, ThermoFisher Scientific) were added to the solutions according to the manufacturer instructions and the incubation was continued for another 16 hours. The beads were washed three times according to the manufacturer’s instructions. Both the beads and the incubation media were measured using a 2250-CA Tri-Carb liquid scintillation analyzer (Packard, USA). Beads counts were normalized to total counts (media and beads counts). Sources and concentrations of the apoB antibody used are summarized in the Major Resources Table.

For quantification of apoB secretion by ELISA, Huh-7 cells transfected with siRNAs and processed as previously described previously^23^ ApoB concentrations in the harvested media were determined by using commercially available ELISA kits (apoB:cat. ELH-ApoB Raybiotech, Norcross, USA), according to the manufacturer’s instructions.

### Quantification of cellular sterols, cholesteryl esters and triglycerides

48 hours after transfection of Huh7 cells with siRNAs, the medium was replaced with either DMEM supplemented with 0.5% FBS, 1% P/S for the measurement of sterols or with DMEM without phenol red (cat. D1145, Sigma-Aldrich) for the measurement of cholesteryl esters and triglycerides. 24 hours later the cells were harvested using Accutase (cat. A6964, Sigma-Aldrich), centrifuged at 1,200 RCF for 5 minutes at 4°C and were either directly pelleted and frozen (cellular sterol quantification) or counted using Vi-Cell XR cell viability analyzer (cat. 731196, Beckman Coulter) (quantification of cholesteryl esters and triglycerides) before freezing at -20°C.

The methods for extraction and measurement of sterols by gas chromatography-mass spectrometry (GC-MS) were described previously^23^.

For the measurement of cholesteryl esters and triglycerides by liquid-chromatography – tandem mass spectrometry (LC-MS/MS), the frozen cell pellets were resuspended in 50 μL PBS and extracted with 1 mL Methanol/MTBE/Chloroform (MMC) (4/3/3) containing SPLASH standard mix (cat. 3300707, Avanti polar lipids, Alabama, USA) containing DAG (15:0/18:1-D7), TAG (15:0/18:1(D7)/15:0) and Cholesterol ester (C18:1(D7)) as internal lipid standards. Lipids were extracted at 37°C (1,400 rpm, 60 minutes) using a thermomixer C (Eppendorf, Hamburg, Germany). The single-phase supernatant was collected after centrifugation at 16000 g (10 minutes, RT), dried under N_2_ and frozen at -20°C until MS analysis. For analysis, the lipid pellets were dissolved in 100 μL methanol on a Thermomixer (550 rpm, 1 hour). Extracted lysates were then transferred to MS vials after centrifugation (16,000 g, 10 minutes at RT). Untargeted lipid analysis was performed on a high-resolution Q-Exactive MS analyser (Thermo Scientific) after lipids were separated by liquid chromatography carried out as previously described^43^ with some modifications. Lipids were separated using ACQUITY UPLC CSH C18 Column (150 mm x 2.1 mm, 1.7 µm particle size, Waters) and a Transcend UHPLC pump (Thermo Fisher Scientific). Liquid chromatography was performed with solvents, acetonitrile:water (6:4) with 10 mM ammonium acetate and 0.1 % formic acid, B) isopropanol:acetonitrile (9:1) with 10 mM ammonium acetate and 0.1 % formic acid at a flow rate of 0.260 mL/min. MS2 fragmentation was based on data dependent acquisition (DDA).

### Statistics

All experimental data were analyzed using GraphPad Prism version 10.6. All results with n<5 were analyzed with non-parametric tests, namely Mann-Whitney test for one to one comparisons or Kruskal-Wallis test coupled with Dunn’s test for multiple comparisons when comparing several conditions. Results with n≥5 were first evaluated for normal distribution of all depicted conditions of each experiment using Shapiro-Wilk normality test. If all examined conditions were normally distributed, the data were analyzed with parametric tests, namely unpaired t-test and 1-way ANOVA coupled with Dunnett’s test for multiple comparisons when comparing several conditions. In case of not normal distribution even for a single condition within one experiment, the data were analyzed with non-parametric tests, as described above. For graphs in which all depicted conditions were compared to the respective control, only p values lower than the threshold limit as mentioned in each legend are depicted. All p values were rounded to the third significant decimal digit. The numbers of experiments, the used statistical tests, and, if applicable, adjustments for multiple testing are described in the legends of the respective figures and tables.

### Ethics

For the data in UK10K FH, LIPIGEN, and LURIC, all consents and local review board approvals were in accordance with the respective ethical frameworks as reported^28–30^. The use of clinical data and samples from patients with the COPA syndrome or the COPG1 syndrome was approved by the Institutional Review Boards (IRB) for the protection of human subjects of the University of California in San Francisco (UCSF, IRB protocol 10-02467) and Boston Children’s Hospital (IRB protocol *04-09-113R*). All participants provided written informed consent.

The sampling of blood and livers from the mutant *Copa* and *Copg1* mice were performed as approved by the Institutional Animal Care and Use Committees of UCSF in San Francisco (Mouse protocol number and approval number 202539) and Boston Children’s Hospital (Mouse protocol number and approval number 00001617), respectively. The liver specific knock-out of *Copg1* by the CRISPR-/cas9 technology as well as the sampling of blood and liver were approved by The Central Authority for Scientific Procedures on Animals (CCD; application# AVD10500202115553), as well as by the Animal Welfare Body (IvD) of the University of Groningen (Groningen, the Netherlands).

## Results

### COPI components limit LDL uptake by Huh-7 hepatocarcinoma cells

In our previous image-based genome wide siRNA screen in Huh-7 hepatocarcinoma cells, the knockdown of three out of seven genes encoding subunits of the COPI coatomer complex, namely *COPA*, *COPB1*, and *COPB2,* significantly reduced the uptake of LDL^11^. To validate these findings, we utilized pooled siRNAs from a different vendor than the siRNA screening library to silence these, as well as the genes encoding the remaining subunits of COPI, namely *ARCN1*, *COPG1*, *COPZ1* and *COPE*, as well as *COPG2* and *COPZ2*, which are paralogues of *COPG1* and *COPZ1*, respectively. The knockdowns suppressed the mRNA expression of the targeted genes almost completely (supplemental figure S1A) and the encoded proteins by at least 80% (solid arrows in supplemental figure S2). Of note and as discussed previously^23^, the knockdown also reduced the abundance of some COPI components of the same subunit (hatched arrows and bars of the same colour in supplemental figure S2) without altering their mRNA expression (supplemental figure S1A). We next used flow cytometry to investigate the effects of siRNA interference against each COPI component on the uptake of LDL, whose protein moiety was labelled with Atto-655. Upon knockdown of the indispensable components of the COPI coatomer (*COPA, COPB1, COPB2, ARCN1, COPG1,* or *COPZ1*) the uptakes of Atto655-LDL (figure 1A) were decreased by at least 95% and hence to an even greater extent as upon knockdown of the positive control gene *LDLR* (-79%). Conversely, the knockdowns of *COPE*, which encodes for the thermosensitive and accessory ε-COP subunit^44,45^, as well as *COPG2 or COPZ2,* whose gene products are paralogues of γ-COP and ζ-COP but are not components of the COPI coatomer^46,47^, did not alter the uptake of LDL (figure 1A). The statistical comparison of the summarized data on loss of indispensable COPI subunits (depicted as red bars) versus those on the non-essential COPI genes *COPE*, *COPG2* and *COPZ2* (depicted as blue bars) revealed a significant difference in LDL uptake (figure 1B, p < 0.001). Taken together, the loss of indispensable subunits of the COPI coatomer compromises the uptake of LDL by Huh-7 cells.

### Silencing of COPI genes alters the glycosylation and cellular localization of LDLR

We next investigated whether the absence of COPI coatomer genes affects LDLR expression or function. Silencing of indispensable COPI genes did not alter the mRNA expression of *LDLR* (supplemental figure S1B and S1C) but the migration pattern of anti-LDLR immunoreactive bands (figure 1C). Instead of the typical ∼160 kDa band corresponding to the mature form of the receptor^6^, an indistinct anti-LDLR immunoreactive signal appeared at a lower apparent molecular weight (figure 1C). Deglycosylation with PNGase F or neuraminidase altered the banding pattern of LDLR but not in the same manner as the knockdown of COPI genes, perhaps because of more complex alterations in the glycosylation of LDLR (figure 1D). The silencing of indispensable *COPA*, *COPB1*, *COPB2*, *ARCN1*, *COPG1*, or *COPZ1* but not dispensable *COPE*, *COPG2*, or *COPZ2* also led to the near absence of LDLR at the cell surface as shown by flow cytometry with an antibody against the extracellular domain of LDLR (figure 1E). The comparison of the combined data on indispensable vs. dispensable COPI genes revealed a significant decrease in the cell surface abundance of (p<0.001, figure 1F). To track the receptor’s intracellular fate in the absence of COPI genes, we overexpressed LDLR with a GFP tag attached to its extracellular, amino terminal end in Huh-7 cells. Western blot analysis with either an anti-LDLR or anti-GFP antibody, or flow cytometry analysis measuring the immunoreactivity of GFP on the cell surface, or just the overall emission of the GFP signal, confirmed the successful overexpression of the construct (supplemental figures S3A-S3D). Silencing of *COPG1* in cells overexpressing GFP-LDLR decreased the cell surface levels of both LDLR (figure 2A) and GFP (figure 2B) but markedly increased the overall GFP signal (figure 2C). These data suggest that upon silencing of *COPG1*, a considerable proportion of LDLR is retained intracellularly rather than reaching the cell surface. Indeed, as shown by immunofluorescence microscopy, interference with *COPG1* led to the disappearance of GFP-LDR from the cell surface (figure 2D, supplemental figure S4A) but appearance of larger and condensed GFP-LDLR containing vesicles, which colocalize with GM130-positive Golgi as well as with LAMP1-positive late endosomes/lysosomes (figures 2E, 2F and supplemental figures S4B, S4C) but not with the SEC61A1-positive ER (figure 2G, supplemental figure S4D).

Taken together, the lack of COPI coatomer subunits interferes with the glycosylation and intracellular trafficking of LDLR. The resulting decrease in cell surface abundance explains the reduced uptake of LDL by Huh-7 cells.

### Genetic variants of COPI genes are associated with elevated LDLC

To investigate the impact of COPI gene variants on plasma levels of LDL in humans, we first explored the associations of 5,569 SNPs in the nine COPI genes with plasma lipids in the aggregated data sets of the Global Lipids Genetics Consortium, which encompasses genotype data from >1.65 million individuals^21^. At the Bonferroni adjusted GWAS threshold of p = 9.60*10^−10^, and in line with the reduced LDL uptake into Huh-7 cells lacking *ARCN1*, 48 SNPs of *ARCN1* were associated with significantly higher levels of LDL-C (figure 3A). 40 of the SNPs are in strong or complete linkage disequilibrium (LD > 0.9). Among them, the intronic variant rs17122278 which has a minor allele frequency (MAF) of 0.146, has the strongest association with LDL-C (METAL effect size 0.0187 mmol/L = 0.72 mg/dL per allele, p= 4.74*10^−22^, figure 3A). In the Genotype Tissue Expression (GTEx) data bank^24^, rs17122278 is associated with changes in the expression of several genes in the locus including ARCN1in several organs, for example in full blood (β = -0.12, p = 1.26×10^−8^, figure 3B).

Our analysis of the Whole Genome Sequencing (WGS) data of nearly 500’000 individuals in the UK Biobank (https://www.azphewas.com: Dataset: UK Biobank 500k WGS (v2) Public) ^22^ did not identify any COPI gene variant significantly associated with LDL-C or apoB levels at the threshold p < 10^−8^. We also analyzed WES data from individuals with familial hypercholesterolemia (FH) or LDL-C levels > 190 mg/dL who had no pathogenic variant in the canonical FH genes *APOB*, *LDLR*, or *PCSK9*. Among 71 patients of the UK10K cohort^28^ and 11 patients of the UNIMORE-FH LIPIGEN cohort^29^, three rare missense variants *COPA* (p.R1178C), *COPA* (p.R1083L), and *COPG1* (p.H60L) were identified. All three variants were predicted to be likely damaging by at least one bioinformatic algorithm and more frequent among the FH patients than expected from the data in GnomADv4.1.0^31,32^ (p = 0.00255 or less,

Fisher’s exact test) and UK Biobank ^33^ (p = 0.00174 or less, Fisher’s exact test) (table 1). Six additional likely damaging rare variants, one in *COPA* and five in *COPG1*, were identified by WES in 92 participants of the LURIC study^30^, who underwent coronary angiography and had LDL-C levels > 190 mg/dL adjusted for lipid-lowering treatment (table2). With a summarized prevalence of 6.52%, *COPA* and *COPG1* variants were more frequent in this cohort of hypercholesterolemic CVD patients compared to 1.53*10^−3^ and 2.06*10^−3^ in GnomADv4.1.0^31,32^ (p < 10^−8^, Fisher’s exact test) and UK Biobank^33^ (p = 5.0 * 10^−8^ , Fisher’s exact test), respectively (table 2).

Finally, we investigated the plasma lipids of five American patients heterozygous for pathogenic variants in *COPA* causing a type I interferonopathy^18^ (figure 3C) and four children homozygous for the pathogenic variant p.K652E in *COPG1* causing combined immunodeficiency as well as their heterozygous parents^17^ (figure 3D). Compared to young adults from the NHANES 2008-2013 population^25^, the LDL-C levels of the COPA syndrome patients were below average (figure 3C) except for one female carrier of the COPA(p.E241K) variant who presented with LDL-C levels above average (139 mg/dL vs. 110 mg/dL). Because of their origin from Oman, we compared the plasma lipoprotein levels of the homozygous *COPG1* variant carriers and their heterozygous parents with those of populations from the United Arab Emirates^26,27^. LDL-C levels of affected children were rather low while those of their heterozygous parents were within the normal ranges for the respective control populations (figure 3D).

In summary, the associations of common *ARCN1* variants with both reduced ARCN1 expression and higher plasma levels of LDL-C in the GLGC databank as well as the increased prevalence of several rare *COPA* and *COPG1* variants in hypercholesterolemic populations but not the normal LDL-C levels in patients with COPA- or COPG1-syndromes support the regulatory relevance of COPI for LDLR.

### Altered plasma levels of lipoproteins in mouse models of COPA or COPG1 syndromes as well as in mice with hepatic *Copg1* depletion

To shed more light on the effects of missense mutations and loss of COPI genes on lipoprotein metabolism, we analyzed mice with heterozygosity for the *Copa^E^*^241^*^K^* (*Copa^E^*^241^*^K/+^*) mutation or homozygosity for the *Copg1^K^*^652^*^E^* mutation (*Copg1^K^*^652^*^E/ K^*^652^*^E^*), which are animal models of the COPA and COPG1 syndromes described above^17,36^. In addition, we generated a mouse model with liver-specific ablation of *Copg1* (table 3).

**Table 3:** Concentrations of lipids and activity of transaminases in plasmas of *Copa*^E241K/+^, *Copg1^K^*^652^*^E/K^*^652^*^E^* or *Copg1*^LKD^ and the respective control mice.

|  | *Copa* mutant mice<sup>36</sup> |  |  | *Copg1* mutant mice<sup>17</sup> |  |  | *Liver specific Copg1* knockout down mice |  |  |
| --- | --- | --- | --- | --- | --- | --- | --- | --- | --- |
|  | *Copa*<sup>+/+</sup><br>(n=6) | *Copa*<sup>E241K</sup><br>/+ (n=6) | p<br>value | *Copg1*<sup>+/+</sup><br>(n=10) | *Copg1*<sup>K652E/K</sup><br>652E (n=10) | p<br>value | *EV*<br>(n=9) | *Copg1*<sup>LKD</sup><br>(n=9) | p<br>value |
| **TC**<br>**(mg/dL)** | 105.054<br>±<br>22.867 | 70.895<br>±<br>18.247 | **0.006**<br># | 79.660<br>±<br>12.256 | 64.579<br>±<br>19.552 | **0.045**<br># | 90.230<br>±<br>3.867 | 90.660<br>±<br>11.619 | 0.917* |
| **HDL-C**<br>**(mg/dL)** | 83.205<br>±<br>16.181 | 51.302<br>±<br>17.965 | **0.009** * | 67.015<br>±<br>11.970 | 53.597<br>±<br>18.077 | **0.05**# | 84.258<br>±<br>3.368 | 80.133<br>±<br>10.363 | 0.273* |
| **nonHDL-C**<br>**(mg/dL)** | 22.558<br>±<br>8.264 | 19.335<br>±<br>4.891 | 0.524<br># | 12.645<br>±<br>2.233 | 10.982<br>±<br>3.051 | 0.181* | 6.445<br>±<br>1.934 | 10.312<br>±<br>1.934 | **0.003**# |
| **TG**<br>**(mg/dL)** | 74.546<br>±<br>20.593 | 50.633<br>±<br>8.202 | **0.025** * | 100.350<br>±<br>38.503 | 57.748<br>±<br>10.846 | **0.003** * | 61.704<br>±<br>9.373 | 41.628<br>±<br>6.352 | **<0.001**<br>* |
| **ALT (U/L)** | 25<br>±<br>5.967 | 23<br>±<br>3.808 | 0.535* | 22<br>±<br>6.07 | 24.2<br>±<br>2.86 | 0.314* | 36.857<br>±<br>3.76 | 55.875<br>±<br>9.64 | **<0.001**<br>* |
| **apoB**<br>**(mg/dL)** | NA | NA | NA | NA | NA | NA | 9.399<br>±<br>1.927 | 13.083<br>±<br>3.258 | **0.010** * |
| **apoA-I**<br>**(mg/dL)** | NA | NA | NA | NA | NA | NA | 48.778<br>±<br>9.530 | 55.996<br>±<br>5.584 | 0.068* |

Mutant *Copa*<sup>E241K/wt</sup> and *Copg1*<sup>K652E/ K652E</sup> mice were described previously<sup>17,36</sup>. Mice with liver-specific *Copg1* knock-down (*Copg1*<sup>LKD</sup>) were generated by CrispR/Cas9-technology as described in the methods section. The values are shown as means ± SD. \*: indicates a statistical analysis using unpaired t-test (2-tailed); # indicates statistical analysis using Mann-Whitney test (2-tailed). NA: Not available

Both the *Copa^E^*^241^*^K/+^* and the *Copg1^K^*^652^*^E/ K^*^652^*^E^* mice had significantly lower plasma concentrations of total and HDL-C as well as triglycerides compared to their wild-type littermate controls, but did not differ by nonHDL-C (table 3). In the pooled plasma of both *Copa^E^*^241^*^K/+^* mice and *Copg1^K^*^652^*^E/ K^*^652^*^E^* mice, mass spectrometry revealed higher concentrations of apoB-derived peptides, respectively, compared to littermate control mice (supplemental table S2). The mRNA expression of none of the COPI genes nor any of the examined genes regulating LDL metabolism (*Ldlr, Hmgcr, Apob*) was significantly altered in *Copa^E^*^241^*^K/+^* liver compared to littermate control (supplemental figure S5). Similarly, α-COP protein levels were not significantly different (supplemental figures S6A, S6B), while apoB was upregulated and LDLR was downregulated in *Copa^E^*^241^*^K/+^* mice (supplemental figures S6A, S6C and S6D, S6E, respectively).

We also disrupted the *Copg1* gene in the liver of male mice by tissue-specific CRISPR/Cas9- mediated genome editing. The expression of *Copg1* mRNA and γ-COP protein were significantly decreased in livers of mice injected with the *Copg1* editing vector *(Copg1^LKD^*) compared to those having received the empty vector (EV-control) (supplemental figures S7A and S8A, S8B, respectively). The residual γ-COP protein may be due to incomplete efficacy of the CRISPR/Cas9 technology or expression by non-parenchymal liver cells. The mRNA expression of other COPI genes or candidate genes regulating the production or removal of lipoproteins was not significantly altered (supplemental figure S7) in the livers of *Copg1^LKD^* with the exception of *Ldlr* that was mildly reduced on the mRNA level (supplemental figure S7J). However, Western Blotting did not reveal any difference in hepatic LDLR or apoB levels between *Copg1^LKD^* and EV control animals (supplemental figures S8C to S8F). Histology did not reveal any differences between livers of the two animal groups (supplemental figures S9A-S9C). Quantification of lipids revealed significantly higher concentrations of triglyceride but normal levels of cholesterol in the liver of *Copg1*^LKD^ mice (supplemental figures S9D and S9E). ALT activity as well as the concentrations of nonHDL-C and apoB, were significantly higher in the plasma of *Copg1^LKD^* mice as compared to EV-controls (table 3). The quantification of apoB peptides by mass spectrometry revealed higher levels of apoB-derived peptides in the pooled plasmas of *Copg1^LKD^* mice as compared to those of EV-control mice. These differences were larger for apoB100-specific peptides than for common peptides of apoB100 and apoB48 (supplemental table S3).

In summary, mice with liver-specific knockdown of *Copg1* but not mice expressing immunopathogenic variants of *COPA* or *COPG1,* presented with elevated plasma levels of non-HDL-C. Moreover, *Copg1* knock-down mice presented with hepatosteatosis.

### Silencing of COPI genes increases the secretion of apoB and intracellular lipid storage of triglycerides and cholesteryl esters

We finally investigated whether differences in the production and secretion of apoB may counteract the effects of disturbed LDL uptake and thereby explain the rather modest increases in LDL-C and apoB and hepatic steatosis of mice with knockdown of *Copg1* as well as the normal levels of LDL-C in mice and humans carrying immunopathogenic *COPA* or *COPG1* variants.

Silencing of the essential COPI genes decreased the expression of *APOB* mRNA (figures 4A and 4B) as well as cellular apoB levels (figures 4C to 4E). Upon statistical comparison of the summarized data on lost indispensable COPI subunits versus knock-down of non-essential COPI genes, the ∼24% and 72% reduction of the apoB mRNA and protein levels, respectively, were statistically significant (figures 4B and 4E, both p < 0.001). Using immunofluorescence microscopy, we did not find the localization of apoB grossly altered in cells lacking *ARCN1* (supplemental figure S10). Using S^35^-methionine/cysteine labeling and quantitative immunoassays, we investigated the effects of silencing COPI genes on the secretion of apoB by Huh-7 cells. Silencing *COPG1* but not the paralogous *COPG2* significantly increased the occurrence in the cell culture medium of proteins that were radiolabeled with S^35^-methionine/cysteine and immunoprecipitated with anti-apoB antibodies (figure 4F, p < 0.001). After knockdowns of *COPA, ARCN1,* or *COPG1* our immunoassay also measured 28% to 65% higher apoB levels in the media of Huh-7 cells compared to the non-coding control siRNA or siRNA against the paralogous *COPG2*. However, these increases were not statistically significant (figures 4G and 4H).

**Figure 4.**
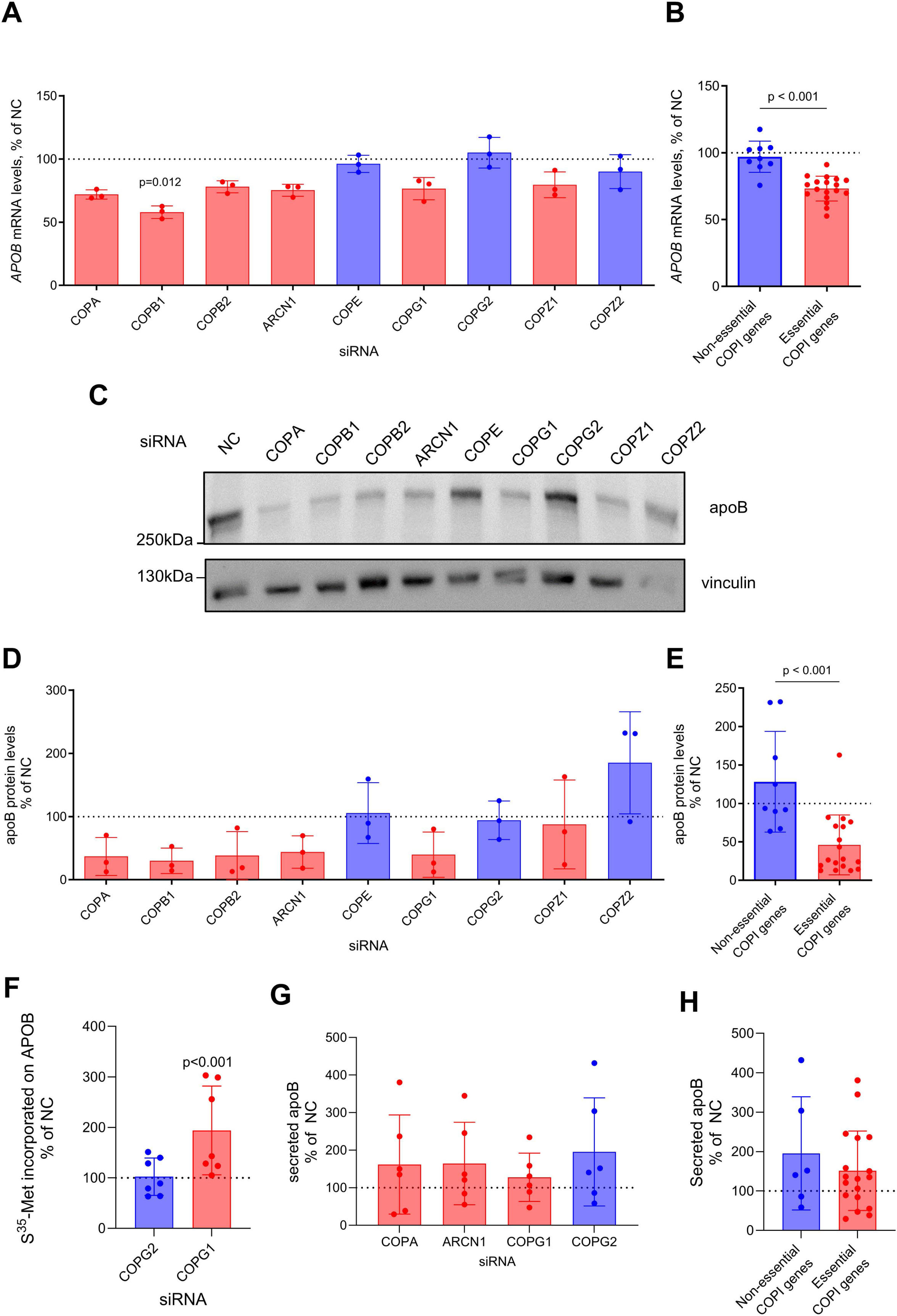
Expression and secretion of apoB by Huh-7 cells depleted of different COPI genes. Wild type Huh-7 cells were transfected with siRNAs targeting the indicated COPI subunits.72 hours after transfection, the cells were either harvested and used for the quantification of *APOB* mRNA levels by qRT-PCR (**A,B**), Western Blotting of cellular apoB protein levels and their respective quantification (**C-E)**), or were treated with methionine and cysteine in 1.5% BSA for 2 hours (**F**). Thereafter, the cells were labelled with 150 μCi/mL of 35S methionine and cysteine. ApoB was immunoprecipitated from the media as described in the methods section. Radioactivity of washed beads was normalized to the total radioactivity in media plus beads as well as the cell number (**F**). Figures **G** and **H** show the quantification of apoB by ELISA in cell cuture media harvested 72 hours after transfection with siRNAs against the the indicated COPI genes. The data shown were normalized to the non-coding control (NC) and are depicted as means ± SD; **C**, **E and H** display merged data on the indispensable COPI subunits (red bars) and the dispensable or paralogous COPI genes (blue bars). Data shown in **A** and **D** are from three independent experiments, while figure parts **F** and **G** display data of seven and six independent experiments, respectively. Statistical analysis of individual experiments was performed using Kruskal-Wallis test with Dunn’s multiple comparisons test between NC and each targeting siRNA (**A, D**) or 1-way ANOVA coupled with Dunnett’s test for multiple comparisons (**F, G**). Pooled data were analysed with either two-tailed unpaired t-test (**B, H**) or two-tailed Mann-Whitney test (**E**). Only statistically significant differences (p<0.05) are indicated.

Mass spectrometry coupled with liquid chromatography (LC-MS) of lysed Huh-7 cells revealed increases of diacylglycerols, triacylglycerols, and cholesteryl esters after knockdown of indispensable COPI subunits which were less pronounced or absent after knockdown of the three non-essential subunits. (figures 5A, 5C and 5E). Compared to the summarized data on knockdowns of non-essential genes, the knockdowns of indispensable COPI genes led to significantly higher cellular levels of these neutral lipids (figures 5B, 5D and 5F, respectively). Gas-chromatography coupled mass spectrometry (GC-MS) of hydrolyzed sterols did not show any significant differences in the cellular content of total cholesterol, phytosterols, or cholesterol precursors between Huh-7 cells expressing or lacking *ARCN1* (supplemental table S4). However, qRT-PCR analysis found the expression of *HMGCR* was slightly increased upon knockdown of indispensable COPI genes, albeit to significantly less degree than upon silencing of non-essential COPI genes (figure 5G and 5H).

**Figure 5.**
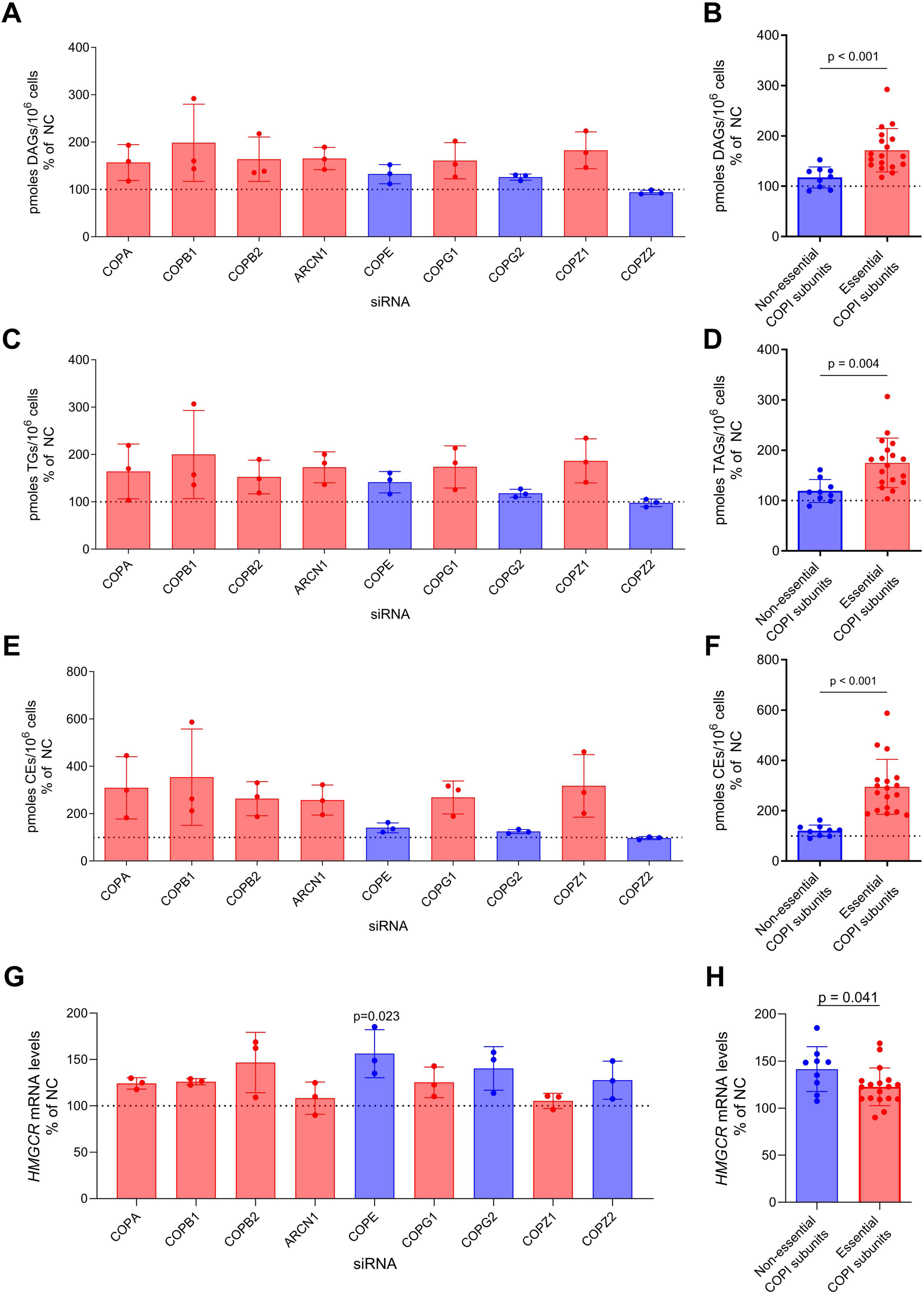
Effect of silencing COPI subunits on cellular diacylglycerols (A, D), triacylglycerols (B, E), and cholesteryl esters (C, F) as well as HMGCR mRNA expression (G, H) by Huh-7 cells. Wild type Huh-7 cells were transfected with siRNAs targeting the indicated COPI subunits. **A**-**F**. 48 hours after transfection, the cells were harvested and treated for 24 hours with DMEM depleted of FBS and phenol red and were collected for the measurement of tri- and di-acylglycerols (TAGs and DAGs, respectively) as well as cholesteryl esters (CEs) by LC-MS. DAGs, TAGs and CEs represent the sum of 12, 97, and 20 species, respectively. **G** and **H**. 72 hours after transfection with the indicated siRNAs, the cells were harvested and used for the quantification of *HMGCR* mRNA levels via qRT-PCR In **A**, **C**, **E**, **G** the data shown were normalized to the NC control and are depicted as means ± SD of 3 independent experiments. **B**, **D**, **F** and **G** display the comparison between the dispensable or paralogous COPI subunits (blue bars) with the non-dispensable COPI subunits (red bars Statistical analyses were performed using either Kruskal-Wallis test coupled with Dunn’s multiple comparison test (**A**, **C**, **E**, **G**), or two-tailed Mann-Whitney test (**B**, **F**) or unpaired two-tailed t-test (**D**, **H**).

Taken together, the lack of COPI coatomer subunits leads to intracellular storage of neutral lipids in Huh7 cells as it does in mouse liver as well as increased ApoB secretion.

## Discussion

In the present study, we provide mechanistic and genetic evidence that the COPI coatomer contributes to the regulation of LDL-C levels in plasma and hepatic lipid storage. By genome-wide RNA interference screening and subsequent validation through targeted gene silencing and flow cytometry, we found that the uptake of LDL into Huh-7 cells is limited by the knock-down of any of the six indispensable components of the COPI coatomer, namely *COPA, COPB1, COPB2, ARCN1, COPG1,* and *COPZ1.* The absent effects of silencing the dispensible *COPE* or the paraologous *COPG2* or *COPZ2* genes corroborate the significance of this finding*: COPE* is the least evolutionary conserved COPI subunit, and its main role is to stabilize COPA^45^. In the mutant CHO cell line ldlF, the lack of *COPE* interferes with the trafficking of cargo at 39.5°C but not at 34°C^44,45^. The 37°C condition of our experiments appears to make COPE dispensable for the functions of the COPI complex, at least for LDL and HDL uptake as shown by us here and previously^23^, respectively, as well as for the regulation of inflammatory cGAS/STING signaling as shown by Steiner and colleagues ^48^. *COPG2* and *COPZ2* are paralogs of *COPG1* and *COPZ1,* respectively. Their functions are not well understood but do not appear to limit the functionality of the COPI coatomer^46,47^.

By contributing to the transport of proteins either from the Golgi apparatus to the endoplasmic reticulum or between the Golgi cisternae, the COPI coatomer plays an important role in the processing and quality control of proteins^13^. With regard to the retrograde Golgi-to-ER transport, it is noteworthy that the SREBP cleavage activating protein SCAP undergoes retrograde transport and thereby fine-tunes the activity of SREBP and thereby the expression of their sterol regulated target genes^49^. However, in our hands the knock-down of COPI genes only led to a mild increase and no change in the mRNA expression of *HMGCR* and *LDLR*, respectively, which are targets of SREBP2. Interestingly, however, we found the expression of *APOB* to be reduced upon loss of indispensable COPI genes.

Our electrophoretic studies indicate limiting effects of COPI function on the glycosylation of LDLR. The band pattern of LDLR upon knockdown of COPI genes was reminiscent of those previously observed in CHO *ldlB*, *ldlC* and *ldlD* mutant cells, which are deficient in *COG1*, *COG2,* and *GALE*, respectively^50^. Together with Golgins and the soluble N-ethylmaleimide-sensitive factor attachment protein receptor (SNARE), the conserved oligomeric Golgi (COG) complex is required for the association of the COPI coat with *cis*-Golgi membranes during intra-Golgi trafficking as well as the delivery of glycosylation enzymes^51^. GALE is a key enzyme of the Leloir galactose metabolic pathway that provides the sugar precursors needed for both N-and O-glycosylation^52^. Of note, our lab also found SR-BI and ABCA1 with aberrant electrophoretic mobility probably due to misglycosylation upon knock-down of COPI genes in Huh7-cells^23^. Moreover, a *de novo* mutation in *ARCN1* was previously identified as the cause of a transient N-glycosylation deficiency during episodes of acute illness^53^. Thus, our results further emphasize the importance of the COPI coatomer in the proper glycosylation of proteins.

The loss of COPI genes diminished the cell surface abundance of LDLR either due to, or in parallel with defective glycosylation. Published data on the role of glycosylation for LDL receptor trafficking and activity are controversial. On the one hand, FH-causing class 2 variants of LDLR contain O-linked carbohydrate chains with core N-acetylgalactosamine residues and N-linked carbohydrate chains of the high mannose type that are only partially trimmed. These variants are properly synthesized and glycosylated in the ER but do not reach the cell surface^3,54^. Moreover, the interference with the attachment of fucose molecules to N- or O-glycans by knockdown of *FUT8* encoding α-1,6-fucosyltransferase promoted degradation of LDLR by induction of PCSK9^55^. On the other hand, patients with type I congenital disorders of N-glycosylation presented with hypobetalipoproteinemia due to increased LDL uptake, which was paralleled with increased expression of SREBP2 protein and cell surface abundance of LDLR^56^. It thus appears that the loss of COPI genes interferes with some glycosylation needed for the proper cell surface trafficking of LDLR.

*In vitro* but not *in vivo*, the knockdown of COPI genes suppressed the expression of *APOB* mRNA or apoB protein. Nevertheless, we found the secretion of apoB from Huh7 cells unaltered as determined by ELISA or even increased if determined by S^35^ methionine labeling. The unaltered or even increased apoB secretion contrasts with the decreased VLDL secretion from McArdle cells^57^ overexpressing a dominant negative mutant of the ADP-ribosylation factor 1 (ARF1). Classically, ARF1 is considered as a limiting factor for the formation of COPI coatomers^58^, but it is increasingly recognized that ARF1 also plays important roles in the regulation of exocytosis, endo-lysosomal trafficking and other coatomer-independent actions. Thus, ARF1 inhibition depletion may compromise apoB trafficking also independently of COPI function. In fact, ARF1 was shown to recruit kinesin motors to triglyceride-rich lipid droplets in the liver and to thereby promote VLDL assembly and secretion^59^.

The COPI coatomer was also shown to recruit both triglyceride degrading and synthesizing enzymes to lipid droplets^60^. In agreement with the finding of disturbed lipolysis upon loss of COPI subunits in Drosophila cells^61^, we observed elevated amounts of triglycerides in both livers of mice with hepatic *Copg1* knock-out and in Huh7 cells after knockdown of indispensable COPI proteins. The coincidence of hepatocellular lipid accumulation with increased apoB secretion and decreased lipoprotein uptake upon loss of COPI function resembles the situation in patients with metabolic syndrome or insulin resistance, who frequently present with fatty liver disease and show both increased apoB production and decreased LDL-catabolism in turnover studies^62^. However, unlike the carriers of COPI variants or *ARCN1* SNPs, these patients present with hypertriglyceridemia. The paradoxical increase of cholesteryl esters despite reduced lipoprotein uptake upon loss of COPI function may be the result of dysregulated intracellular esterification and/or hydrolysis of cholesteryl esters. In fact, defective lysosomal hydrolysis of cholesteryl esters and triacylglycerols in patients with cholesteryl ester storage disease due to acid lysosomal acid lipase deficiency is also accompanied by increased production of apoB containing lipoproteins. However, LDLR activity and LDL catabolism are normal in these patients^63,64^.

By limiting LDLR-mediated LDL uptake, the loss of COPI coatomer function is expected to increase LDL-C levels. In agreement with this hypothesis, our analysis of the GLGC data of more than 1.65 million persons ^21^ identified many SNPs in *ARCN1* to be associated with higher plasma levels of LDL-C. Some of them are associated with reduced *ARCN1* expression and were previously identified in Asian populations to be associated with differences in LDL-C^65,66^. Also, in agreement with the limiting effect of the COPI coatomer on LDLR activity, we found rare coding variants of *COPA* and *COPG1* enriched in hypercholesterolemic subjects who have no causal variant in the classical familial hypercholesterolemia genes *LDLR, APOB, or PCSK9*. Likewise, mice with a CRISPR/Cas9-mediated knock-down of hepatic *Copg1* had significantly higher plasma levels of nonHDL-C and apoB100.

Conversely, patients and mice with the COPA- or COPG1-syndrome had normal plasma levels of LDL-C and nonHDL-C, respectively. Although our analyses are limited by the small sample size of FH cases, the heterogeneous effects of distinct *COPG1* and *COPA* variants on LDL-C or non-HDL-C may reflect different effects of the individual variants on the cellular fate of apoB, LDLR and other proteins contributing to the metabolism of LDL. In fact, depending on the domain structurally altered by mutations causing the COPA syndrome, either only retrograde Golgi-to-ER transport or both retrograde and anterograde ER-to-Golgi transport of stimulator of interferon genes (STING) were found to be defective^16–18^.

In conclusion, the loss of COPI coatomer subunits interferes with the glycosylation and cell surface abundance of LDLR as well as the uptake of LDL and apoB secretion by cultured human hepatocarcinoma cells. The association of common COPI gene variants with higher levels of LDL-C and the enrichment of rare coding variants in hypercholesterolemic patients indicates the physiological relevance of these regulatory steps in humans. However, the rather low LDL-C levels in humans and mice carrying rare immunopathogenic *COPA* and *COPG1* variants indicate more diverse structure-function relationships of the COPI coatomer in LDL metabolism. Notwithstanding this complexity, our data highlight the COPI coatomer as an essential factor in lipid and lipoprotein metabolism and, thereby, as a determinant of plasma lipoprotein concentrations.

## Nonstandard Abbreviations and Acronyms

FH: Familial hypercholesterolemia
RNAi: RNA interference
GWAS: Genome wide association study
WES: Whole exome sequencing
WGS: Whole genome sequencing
CRISPR/Cas9: Clustered regularly interspaced short palindromic repeats/ CRISPR-associated protein 9
GC-MS: Gas chromatography mass spectrometry
LC-MS: Liquid chromatography mass spectrometry
LDL-C: Low-density lipoprotein cholesterol
HDL-C: High-density lipoprotein cholesterol
TG: Triglycerides
FC: Free cholesterol
CE: Cholesteryl ester
apoB: Apolipoprotein B
COPI: Coat protein I
*COPA*: α-COP,COPI coat complex subunit alpha
*COPB1*: β-COP,COPI coat complex subunit beta 1
*COPB2*: β’-COP,COPI coat complex subunit beta 2
*ARCN1*: δ-COP,COPI coat complex subunit delta
*COPE*: ε-COP,COPI coat complex subunit epsilon
*COPG1*: γ-COP,COPI coat complex subunit gamma 1
*COPG2*: γ’-COP,COPI coat complex subunit gamma 2
*COPZ1*: ζ-COP,COPI coat complex subunit zeta 1
*COPZ2*: ζ’-COP,COPI coat complex subunit zeta 2
LDLR: LDL receptor

## AKS is supported by NIH grant R01AI168299

R.G is funded by grant AI139633 - 5R01AI139633-05 from NIH

The research on exon variants in UK Biobank has been conducted using summary statistics generated by GeneBass from the UK Biobank resource (under application 26041 and 48511).”

Flow cytometry was performed with equipment of the flow cytometry facility, University of Zurich.

## Highlights

- RNA interference with any essential components of the COPI coatomer decreases the uptake of LDL holoparticles into Huh-7 hepatocarcinoma cells by interfering with the glycosylation and cell surface abundance of LDLR.
- Silencing of *COPA*, *COPB1*, *COPB2*, *ARCN1*, *COPG1*, and *COPZ1* also promotes apoB secretion and intracellular lipid storage in Huh7 cells.
- Single nucleotide polymorphisms of *ARCN1 are* associated with significantly higher LDL-cholesterol (LDL-C) levels in the population and some rare variants of COPA and COPG1 are enriched in hypercholesterolemic patients
- Liver-specific knock-down of COPG1 leads to higher plasma levels of nonHDL-cholesterol and hepatic triglyceride accumulation in mice.

## Supplemental Material

### 1. Supplemental figures

**Supplemental Figure S1.**
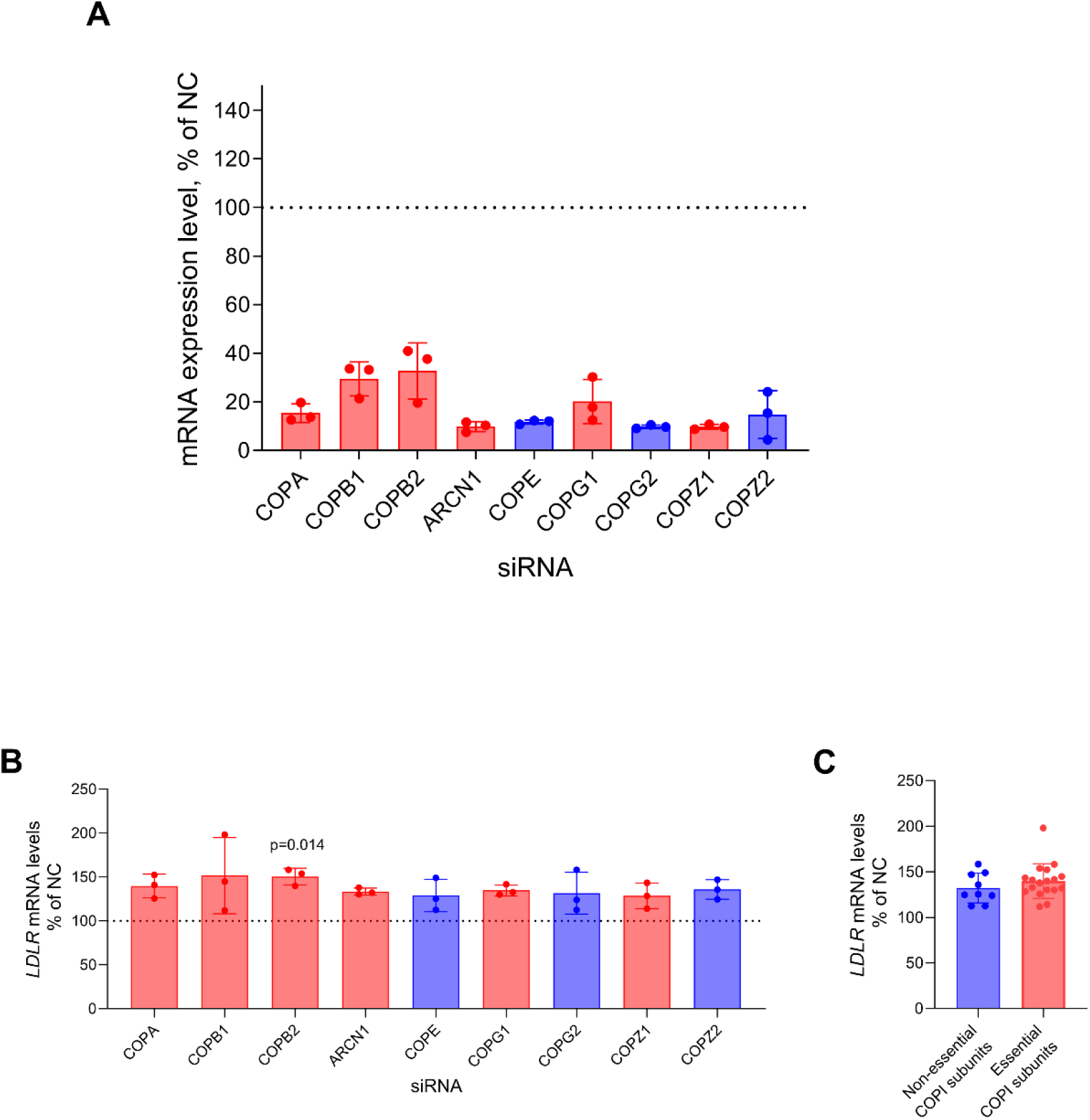
Silencing efficiency and effects of knocking down different COPI genes on LDLR mRNA levels. Huh-7 cells transfected with the indicated siRNAs were collected 72 hours post transfection and used for the quantification of the indicated mRNAs by using qRT-PCR. The data are normalized to the non-coding control (NC = 100%) and are shown as means ± SDs of three independent experiments (**A, B**). **C** compares merged data on the indispensable COPI subunits (red bars) and the dispensable or paralogous COPI genes (blue bars). Statistical analysis was performed using either Kruskal-Wallis test coupled with Dunn’s test for multiple comparisons between the non-coding (NC) control siRNA and each targeting siRNA (**B**) or two tailed Mann-Whitney test (**C**). Only statistically significant differences (p<-0.05) are displayed.

**Supplemental Figure S2.**
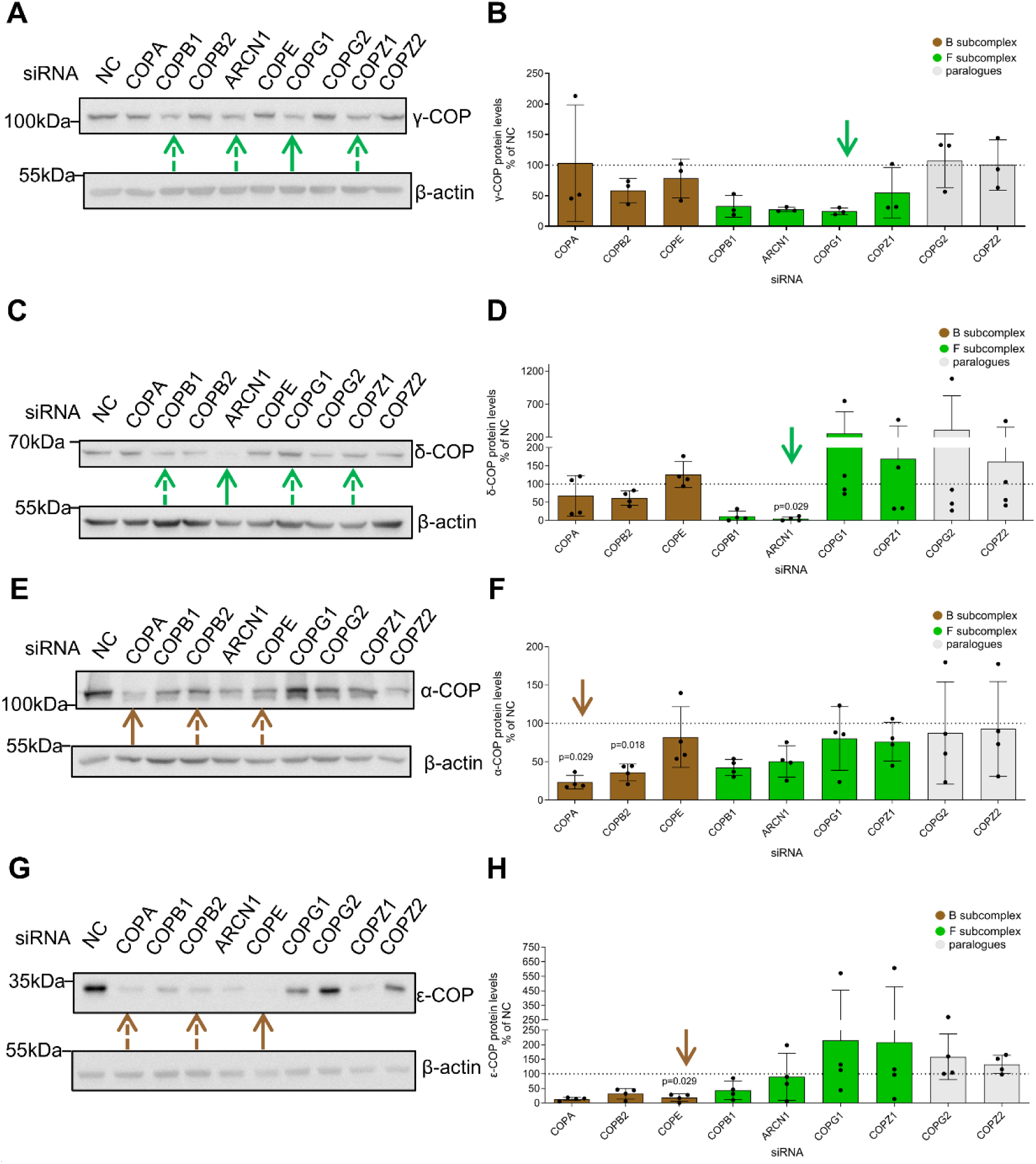
Effects of silencing different COPI subunits on the protein levels of different COPI proteins. Huh-7 cells transfected with the indicated siRNAs were collected 72 hours post transfection and used for immunodetection of γ-COP (**A**), δ-COP (**C**), α-COP (**E**) or ε-COP (**G**) by Western Blotting, with β-actin serving as the loading control. The depicted Western Blots are representative of three (**A**) or four (**C**, **E, G**) independent experiments. Bold arrows indicate the targeted COPI protein directly by siRNA, while the hatched arrows indicate COPI subunits that are part of the same F- (**A**, **C**) or B-subcomplex (**E**, **G**). **B**, **D**, **F** and **H** depict the quantification of all the Western Blot experiments for γ-, δ-, α- and ε-COP subunits. The data were normalized to the non-coding (NC) control condition and are shown as means ± SD of 3 (**B**) or 4 (**D**, **F, H**) experiments. The arrow in each graph indicates the on-target condition. In all cases, statistical analysis was performed using two-tailed Mann-Whitney test between the NC and the on-target condition, and Kruskal-Wallis test with Dunn’s multiple comparisons test between the NC and the remaining conditions. Only statistically significant differences (p < 0.05) are shown.

**Supplemental Figure S3.**
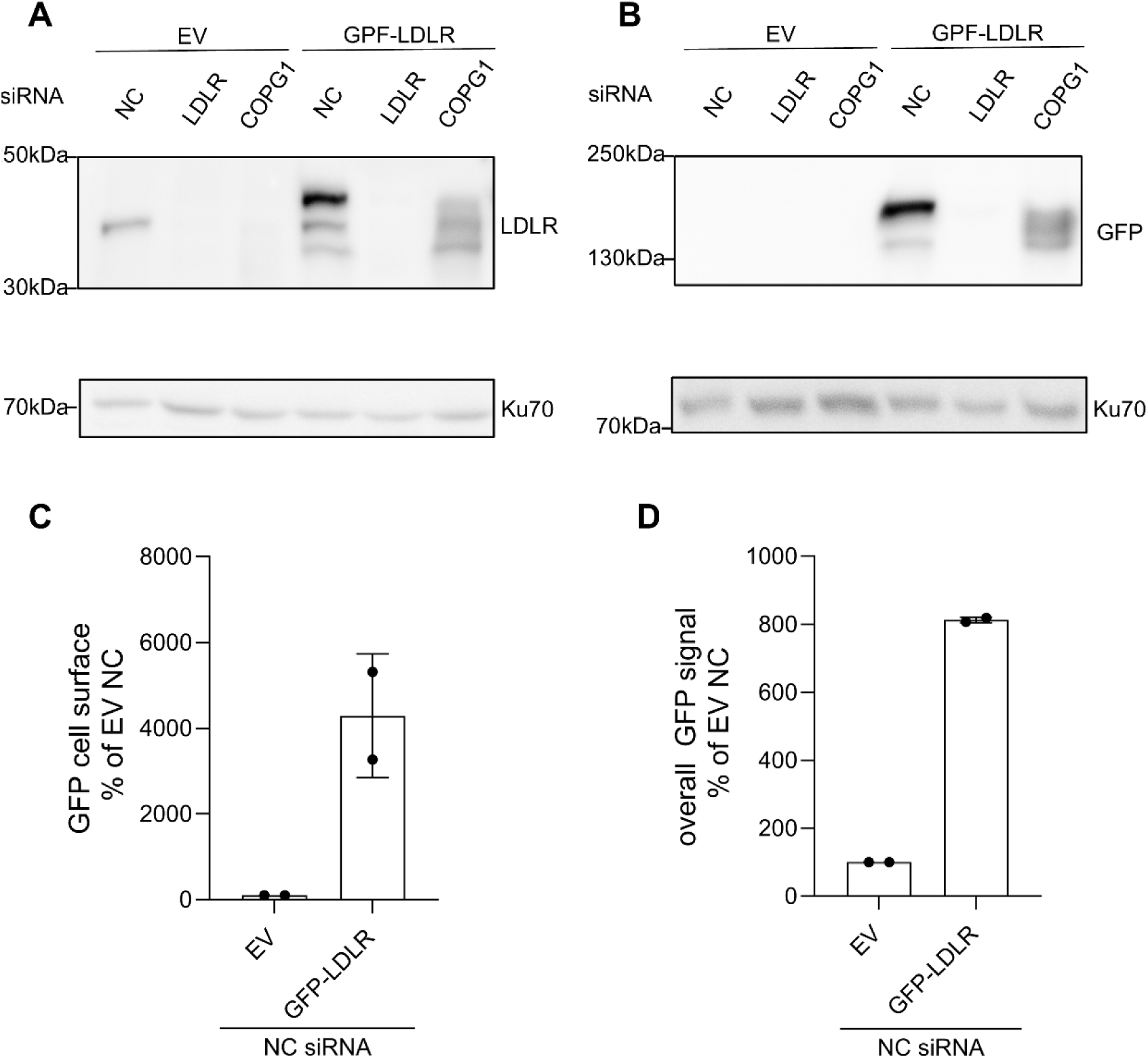
Stable overexpression of GFP-LDLR in Huh-7 cells. Huh-7 cells overexpressing either an empty vector (EV) alone or combined with another vector containing *LDLR* with a GFP tag attached to its aminoterminal end (GFP-LDLR) were transfected with the indicated siRNAs. 72 hours after transfection, the cells were harvested and lysed. Proteins of lysates were electrophoresed and blotted to a membrane for immunodetection of LDLR and GFP with antibodies against the aminoterminal epitope of LDLR (**A**) or GFP (**B**), respectively. In both cases, Ku70 served as the loading control. The Western Blot experiments in **A** and **B** were carried out once. 72 hours after transfection with NC siRNA (**C**, **D**) the cells were collected for the measurement of the GFP cell surface levels using an anti-GFP antibody (**C**) or the entire GFP signal (**D**). The data are normalized to the EV and are shown as means ± SD of two independent experiments.

**Supplemental Figure S4.**
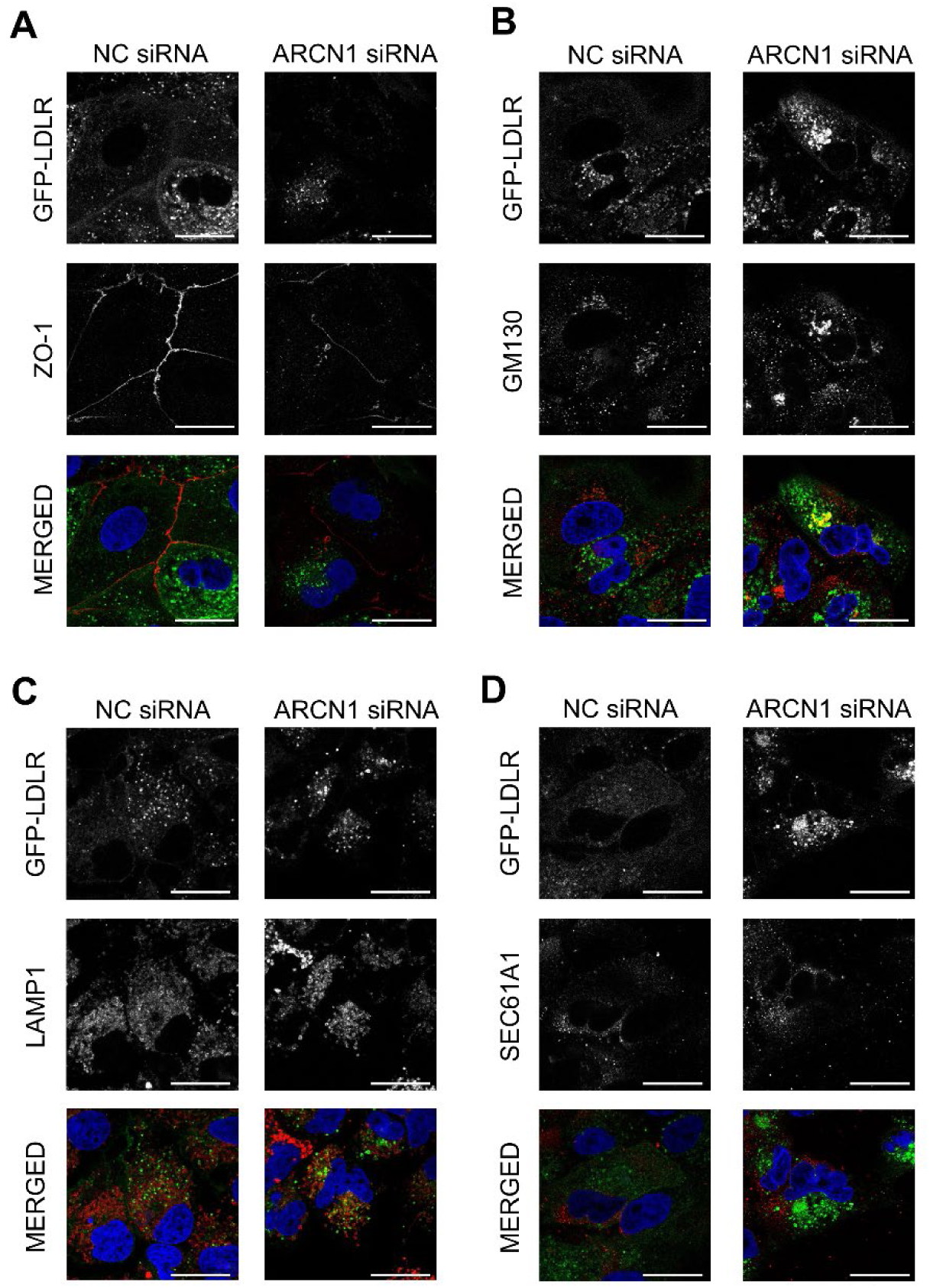
Subcellular localization of LDLR is altered in Huh-7 cells with knock down of ARCN1. Confocal micrographs of GFP-LDLR and ZO-1 (**A**), GM130 (**B**), LAMP1 (**C**) and SEC61A1 (**D**) using Huh-7 cells overexpressing GFP-LDLR 72 hours after transfection with the indicated siRNAs. The micrographs of the merged signals are the ones presented in the Figure 2. The data shown represent two independent experiments. The scale bar for all micrographs is 25 μm.

**Supplemental Figure S5.**
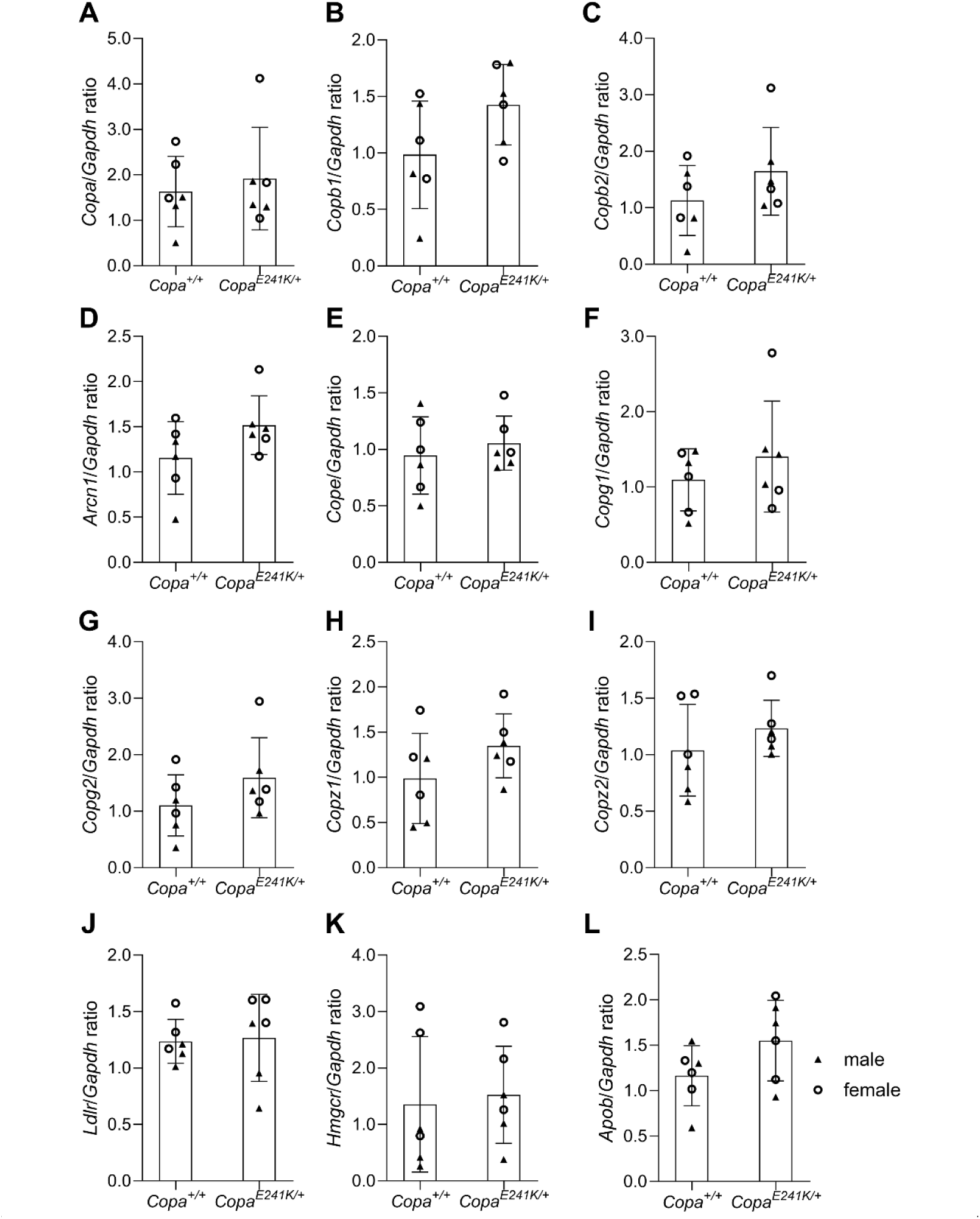
qPCR data of livers from Copa *^E^*^241^*^K/+^* mice. Livers from six *Copa* mutant mice and six wild type littermates were collected, frozen in liquid nitrogen, and shipped on dry ice to be processed for the determination of the mRNA levels of *Copa* (**A**), *Copb1* (**B**), *Copb2* (**C**), *Arcn1* (**D**), *Cope* (**E**), *Copg1* (**F**), *Copg2* (**G**), *Copz1* (**H**), *Copz2* (**I**), *Ldlr* (**J**), *Hmgcr* (**K**) or *Apob* (**L**) via qRT-PCR. For each sample, the ratio of each gene of interest relative to *Gapdh* was calculated. The data are shown as means ± SDs of 6 animals per each genotype. Male and female sexes of each animal are indicated with closed triangles and open circles, respectively. Statistical analyses using two-tailed Mann-Whitney (**A**) or unpaired two-tailed t-tests (**B**-**L**) did not reveal any statistically significant differences between mutant and wild type mice.

**Supplemental Figure S6.**
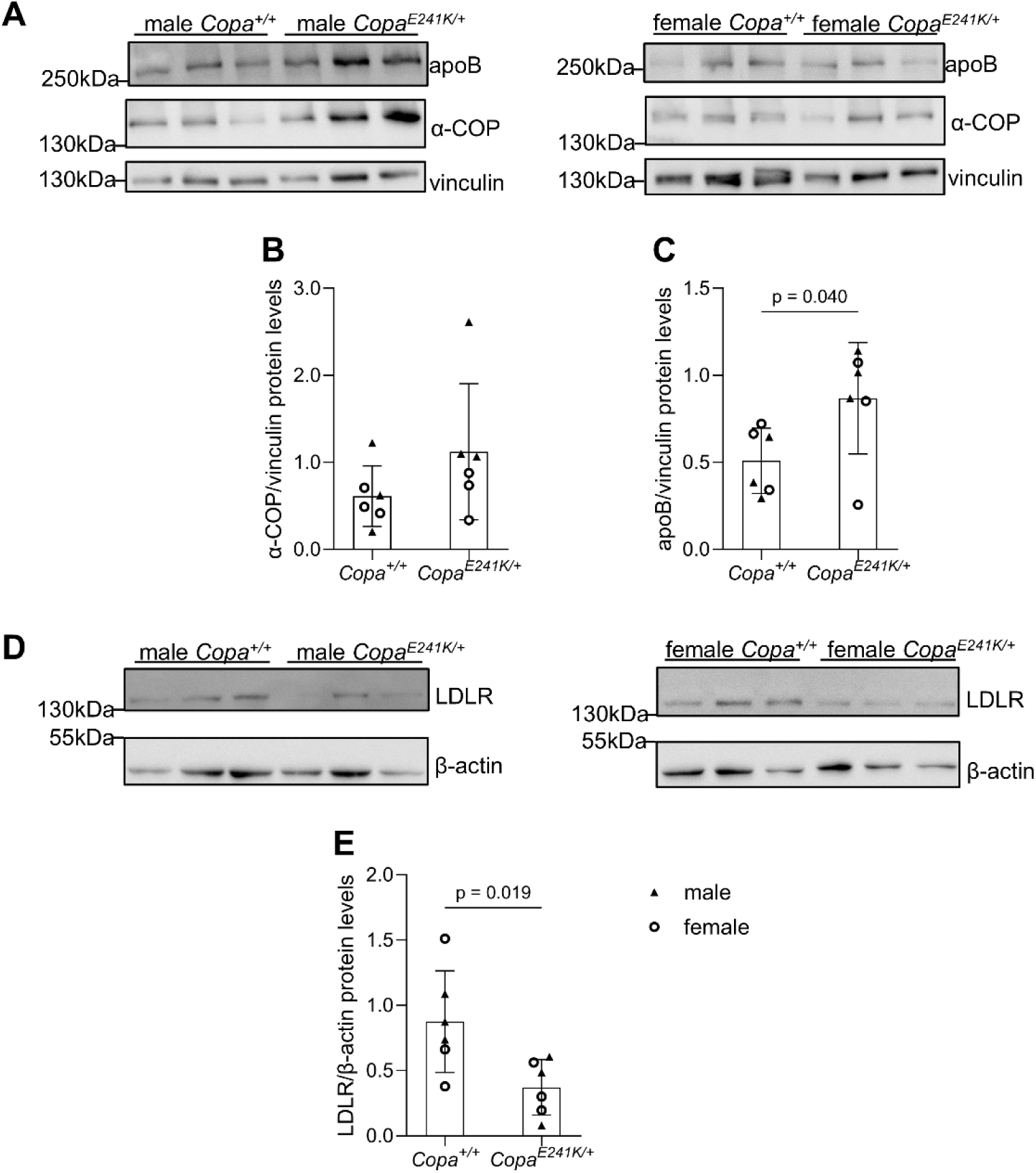
Western Blots of livers from *Copa^E^*^241^*^K/+^* mice. Proteins of homogenized liver lysates from *Copa* mutant mice and their wild type littermates were electrophoresed and electroblotted to PVDF sheets for immunodetection of α-COP (**A**, **B**), apoB (**A**, **C**) and LDLR (**D**, **E)**. For α-COP, apoB and LDLR, the actual western blot image is shown in **A** and **D**, together with graphs of their respective densitometric quantification in **B, C** and **E**, respectively. In all graphs (**B, C, E**), densities of each protein of interest are shown relative to the loading control (**B**, **C**: vinculin; **E**: β-actin). The data are means ± SDs of 6 animals per genotype. Male and female sexes of each animal are indicated with closed triangles and open circles, respectively. Statistical analysis was performed using unpaired two-tailed t-tests (**B**, **C**, **E**), without correction for multiple testing. Only statistically significant comparisons (p<0.05) are shown.

**Supplemental Figure S7.**
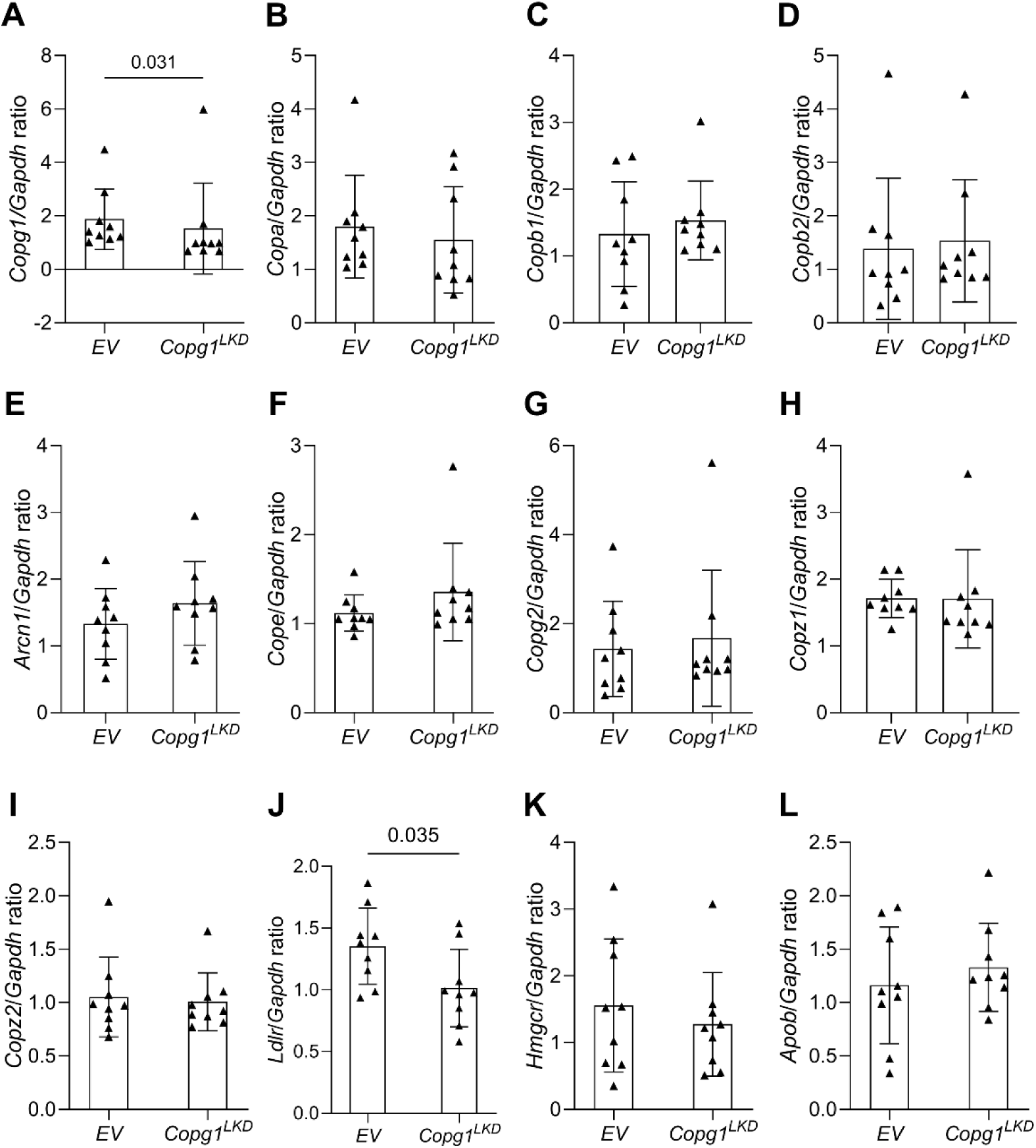
qPCR data of livers from *Copg1^LKD^* mice. Livers from nine *COPG1^LKD^* mice and nine littermates were collected, frozen in liquid nitrogen, and shipped on dry ice to be processed for determination of the mRNA levels of *Copg1* (**A**), *Copa* (**B**), *Copb1* (**C**), *Copb2* (**D**), *Arcn1* (**E**), *Cope* (**F**), *Copg2* (**G**), *Copz1* (**H**), *Copz2* (**I**), *Ldlr* (**J**), *Hmgcr* (**K**), or *Apob* (**L**) via qRT-PCR. For each sample, the Ct-values of each gene of interest relative to those for *Gapdh* were calculated. The data are shown as means ± SDs of nine animals per genotype. Statistical analysis was performed using two-tailed Mann-Whitney (**A-D, F-I**) or unpaired two-tailed t-tests (**E**, **J**-**L**), without any correction for multiple testing. Only statistically significant differences (p<0.05) are shown.

**Supplemental Figure S8.**
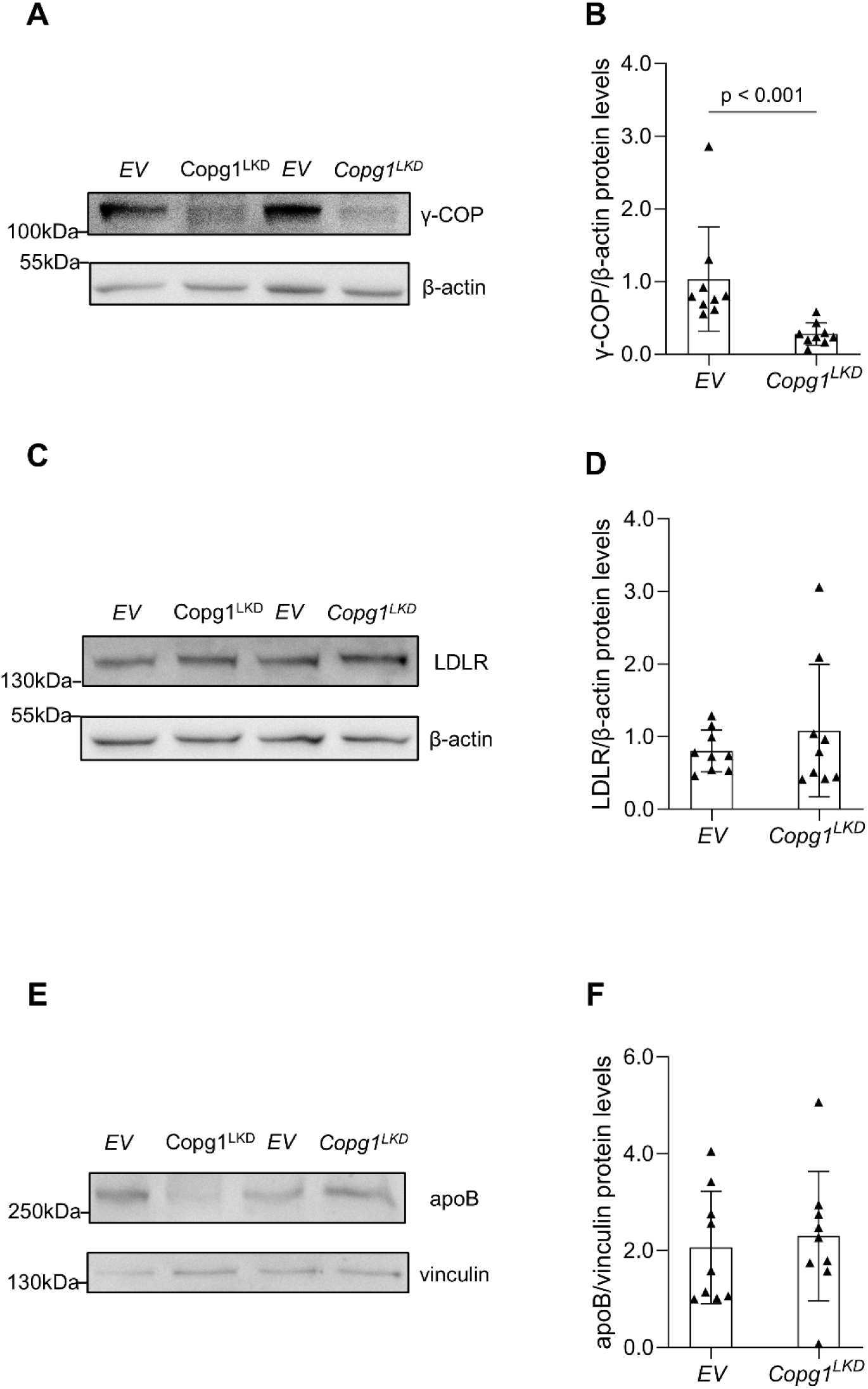
Western Blots of livers from *Copg1^LKD^* mice. Proteins of homogenized liver lysates from *Copg1^LKD^* mice and their littermates were electrophoresed and electroblotted to PVDF sheets for immunodetection of γ-COP (**A**, **B**), LDLR (**C, D**), or apoB (**E, F**). The graphs represent the densities of γ-COP (**B**), LDLR (**D**), or apoB (**F**) relative to the respective loading control (**B**, **D**: β-actin; **F**: vinculin). The data are shown as means ± SDs of nine animals per genotype. Statistical analysis was performed using two-tailed Mann-Whitney (**B, D**) or two-tailed unpaired t-test (**F**) without any correction for multiple testing. Only statistically significant differences (p<0.05) are displayed.

**Supplemental Figure S9.**
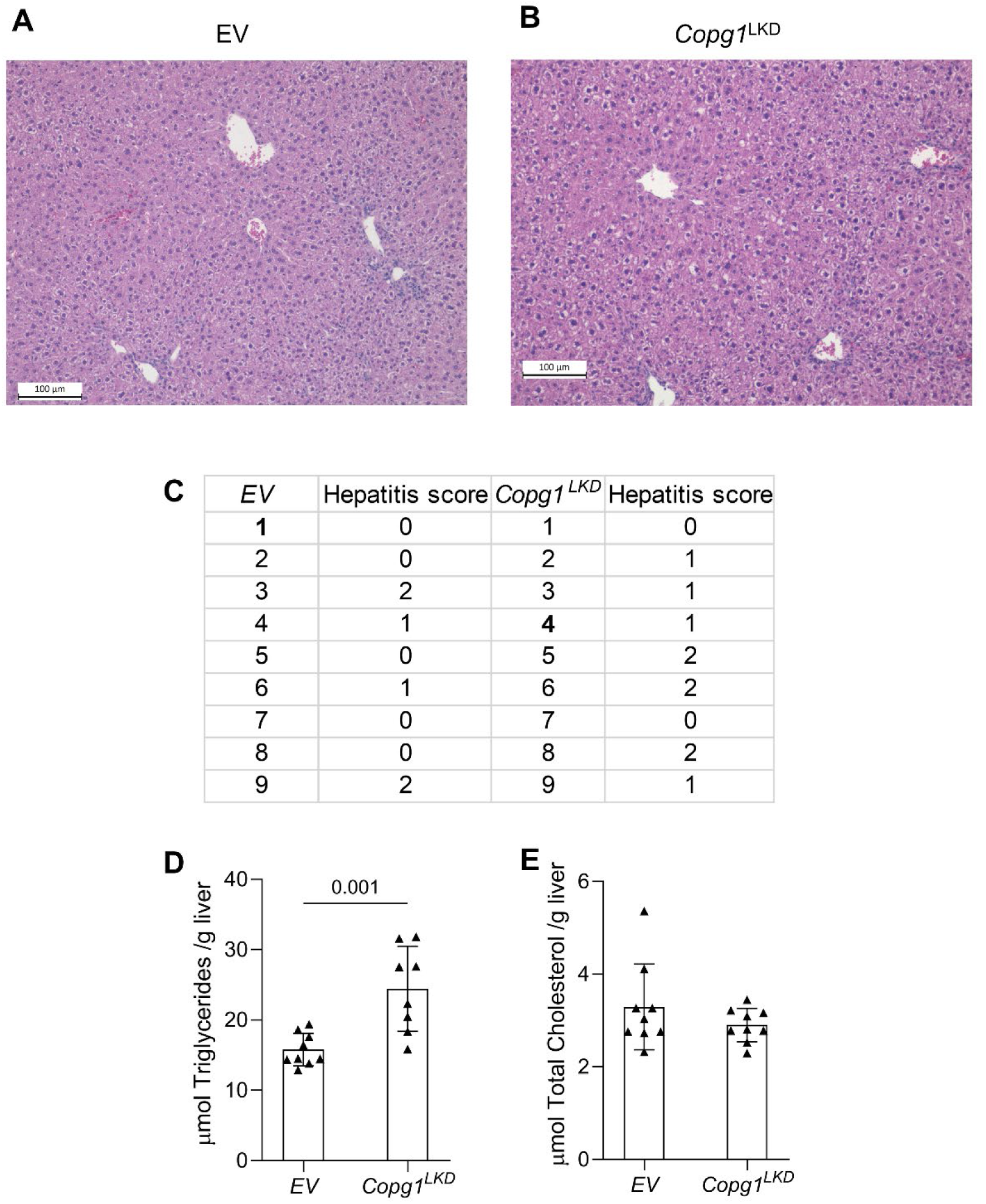
Histology and lipid content of livers from *Copg1^LKD^* mice. Histology (**A, B,** and **C**), triglycerides (**D**) or cholesterol (**E**) were determined in liver sections obtained from *Copg1^LKD^* mice. **A** and **B** show representative liver sections stained with H&E of mice injected with a virus containing sgEV (EV) (**A**) or sgCOPG1 (*Copg1^LKD^*) **(B)**. The table (**C**) shows hepatitis scores for individual animals. Bold letters indicate the animals whose liver sections are shown in **A** and **B**. The bar diagrams show concentrations of triglycerides (**D**) and cholesterol (**E**) per gram of liver tissue. Statistical analysis was performed using two-tailed Mann-Whitney [**D**, p< 0.001, N = 8 (EV), 9 (Copg1^LKD^)] or unpaired two-tailed t-test [(**E,** not significant, N = 9 (both EV and Copg1^LKD^)].

**Supplemental Figure S10.**
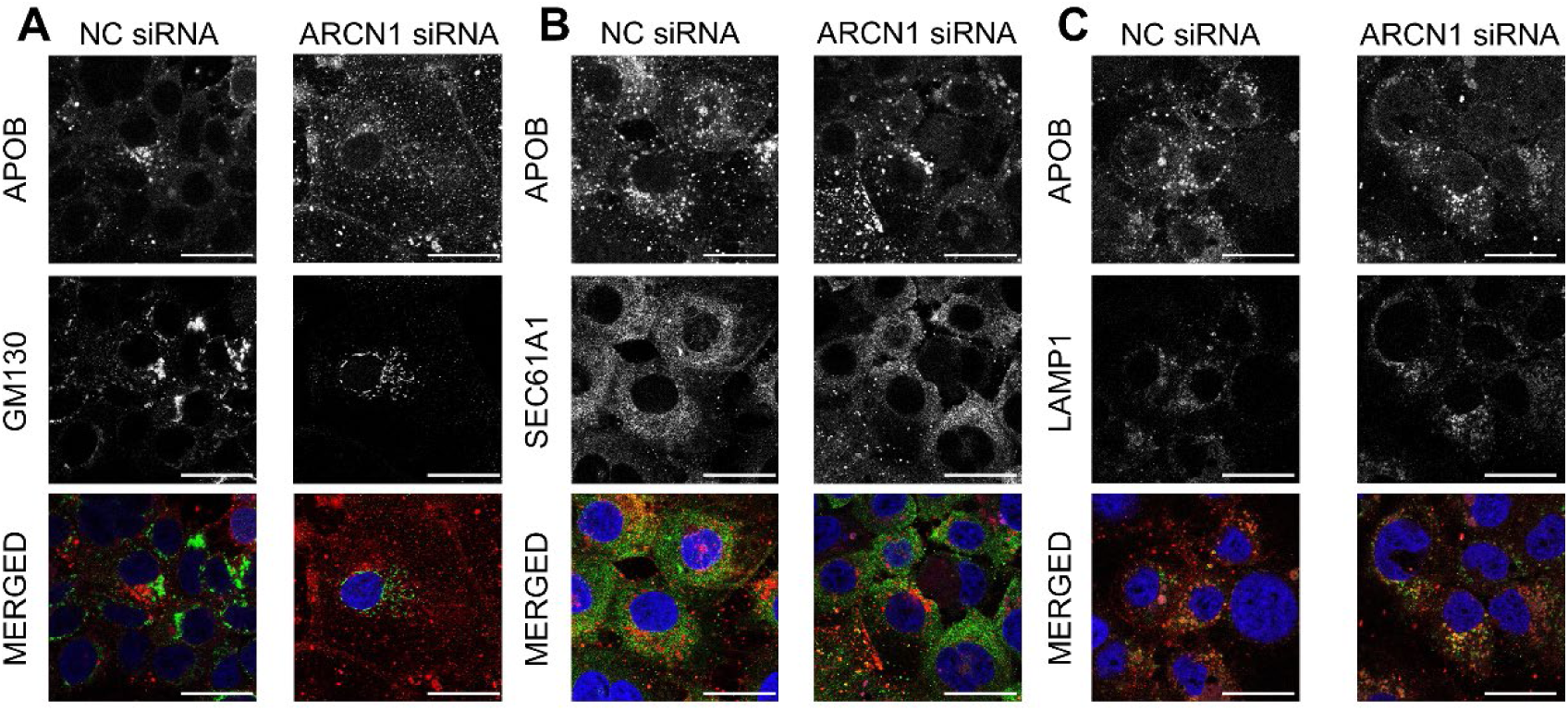
Subcellular localization of apoB in Huh-7 cells depleted of ARCN1. Confocal micrographs of apoB and GM130 (**A**), SEC61A1 (**B**) and LAMP1 (**C**) in Huh-7 cells 72 hours after transfection with the indicated siRNAs. The data shown represent two independent experiments. The scale bar for all micrographs is 25 μm.

### 2. Supplemental tables

**Supplemental Table S1.**
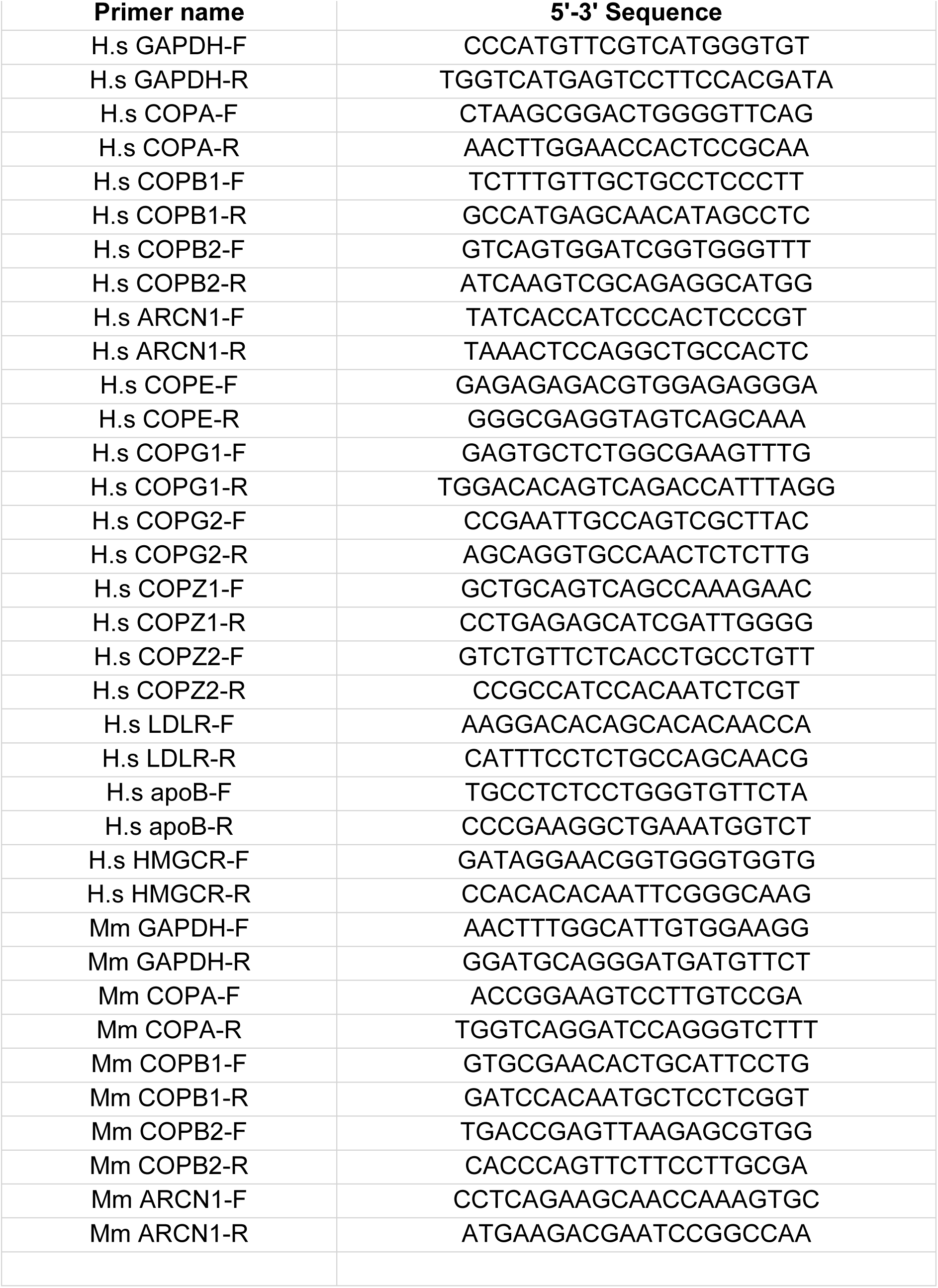
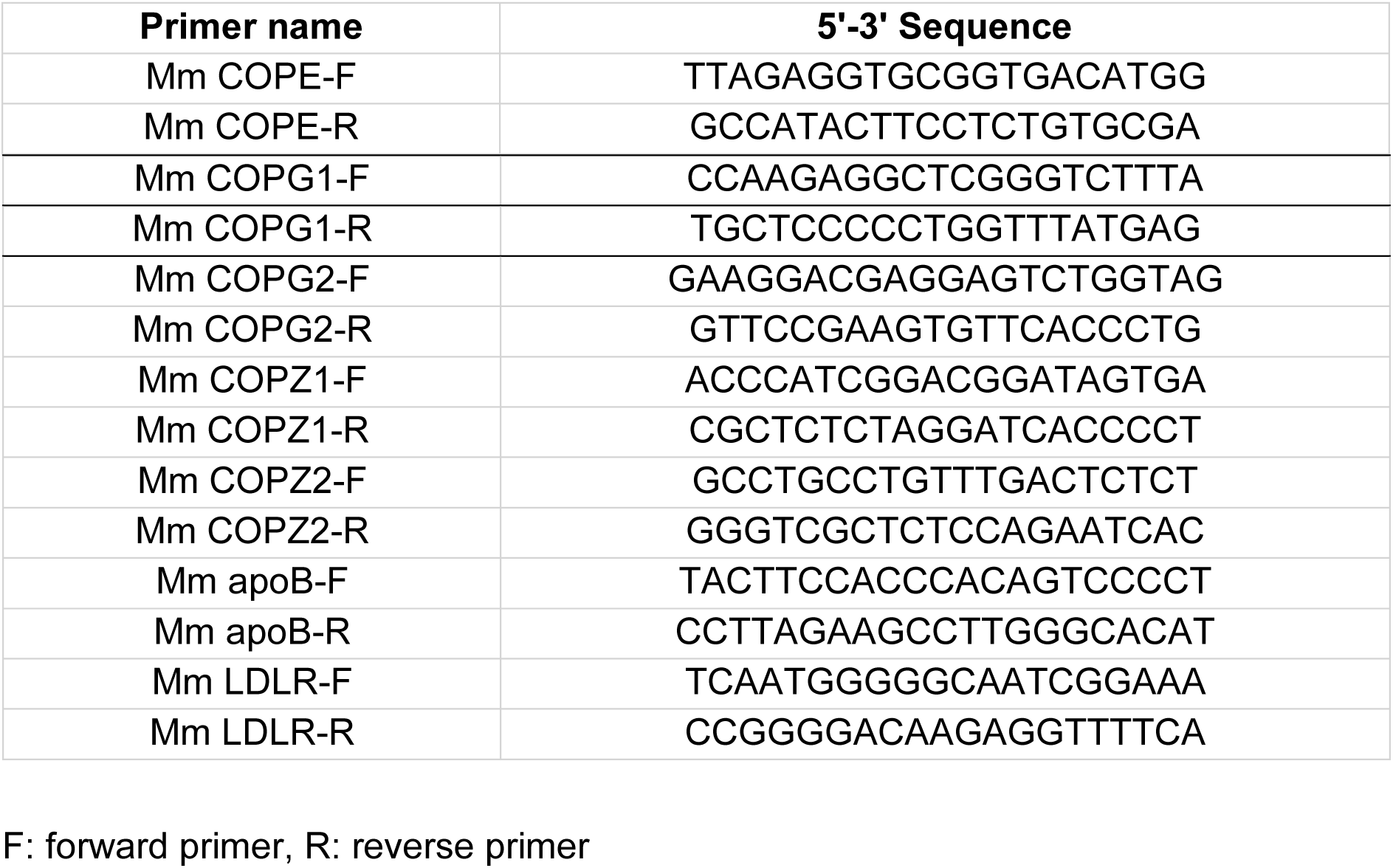
PCR primer sequences for all primer pairs.

**Supplemental Table S2.**
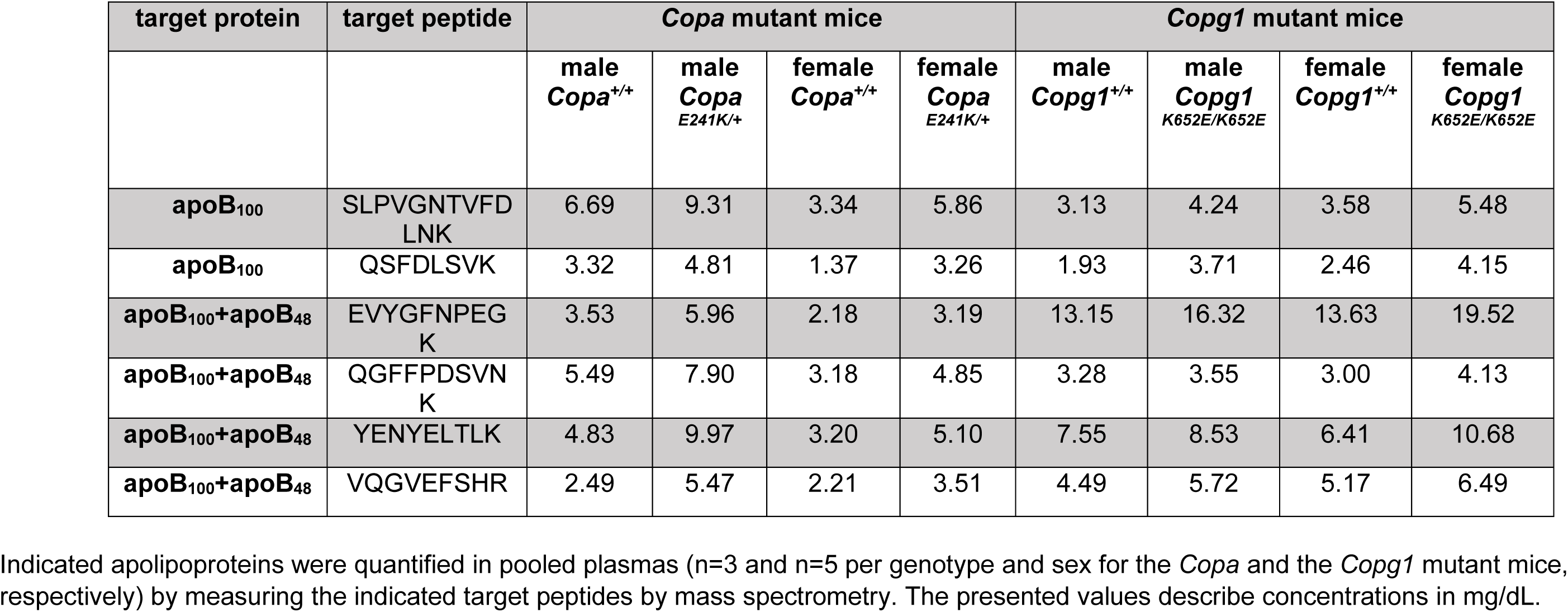
Quantification of apolipoproteins by mass spectrometry plasmas of mice heterozygous for *Copa^E^*^241^*^K^* or homozygous for *Copg1^K^*^652^*^E^* and their respective littermate controls.

**Supplemental Table S3.** Quantification of apolipoproteins by mass spectrometry in *Copg1* knock-down (*Copg1*^LKD^) mice and control (EV) mice.

| target protein | target peptide | EV | *Copg1*<sup>LKD</sup> | p value |
| --- | --- | --- | --- | --- |
| **apoB<sub>100</sub>** | SLPVGNTVFDN<br>K | 1.800 ± 0.579 | 2.226 ± 0.330 | 0.151<sup>#</sup> |
| **apoB<sub>100</sub>** | QSFDLSVK | 1.482 ± 0.509 | 2.050 ± 0.320 | 0.183<sup>#</sup> |
| **apoB<sub>100</sub>+apoB<sub>48</sub>** | EVYGFNPEGK | 9.046 ± 2.187 | 9.438 ± 1.067 | 0.310<sup>#</sup> |
| **apoB<sub>100</sub>+apoB<sub>48</sub>** | QGFFPDSVNK | 2.320 ± 0.528 | 2.726 ± 0.586 | 0.283<sup>*</sup> |
| **apoB<sub>100</sub>+apoB<sub>48</sub>** | YENYELTLK | 6.402 ± 1.377 | 6.592 ± 0.690 | 0.690<sup>#</sup> |
| **apoB<sub>100</sub>+apoB<sub>48</sub>** | VQGVEFSHR | 3.776 ± 0.570 | 4.054 ± 0.497 | 0.435<sup>*</sup> |

Indicated apolipoproteins were quantified in plasmas from mice injected with AAV containing sgEV or sgCopg1 by measuring the indicated target peptides by mass spectrometry. The presented values describe concentrations in mg/dL. The data are shown as means ± SD of 5 measured samples per condition. Statistical analysis was carried out either using 2-tailed unpaired t-test (\*) or 2-tailed Mann-Whitney test (#).

**Supplementary table S4.** Effects of *ARCN1* knockdown on cellular levels of sterols in Huh-7 cells.

| classification | sterol | Basal Condition |  |  |
| --- | --- | --- | --- | --- |
|  |  | NC<br>(ng/mg cell pellet) | ARCN1<br>(ng/mg cell pellet) | p value |
| cholesterol<br>precursors | lathosterol | 2,186±268 | 2,034±68 | 0.700 |
|  | lanosterol | 28.833±3.522 | 15.567±1.950 | 0.100 |
|  | desmosterol | 73.933±6.408 | 80.500±6.421 | 0.400 |
|  | dihydrolanosterol | 10.147±1.181 | 6.450±0.961 | 0.100 |
|  | cholesterol | 37,633±3,355 | 35,067±1,966 | 0.400 |
| phytosterols | stigmasterol | 1.070±0.026 | 1.137±0.091 | 0.700 |
|  | sitosterol | 3.307±0.400 | 2.713±0.202 | 0.200 |
|  | campesterol | 3.083±0.314 | 3.683±0.419 | 0.200 |

The indicated sterols were measured by gas chromatography-mass spectrometry in lipid extracts of pellets from Huh-7 cells which were transfected with siRNAs targeting *ARCN1* or non-targeting siRNAs (NC) and incubated with DMEM+0.5% FBS
The data are shown as mean ± SD of 3 independent experiments. The p values presented are calculated with Mann-Whitney test (two-tailed) and not adjusted for multiple testing.

## Major Resources Table

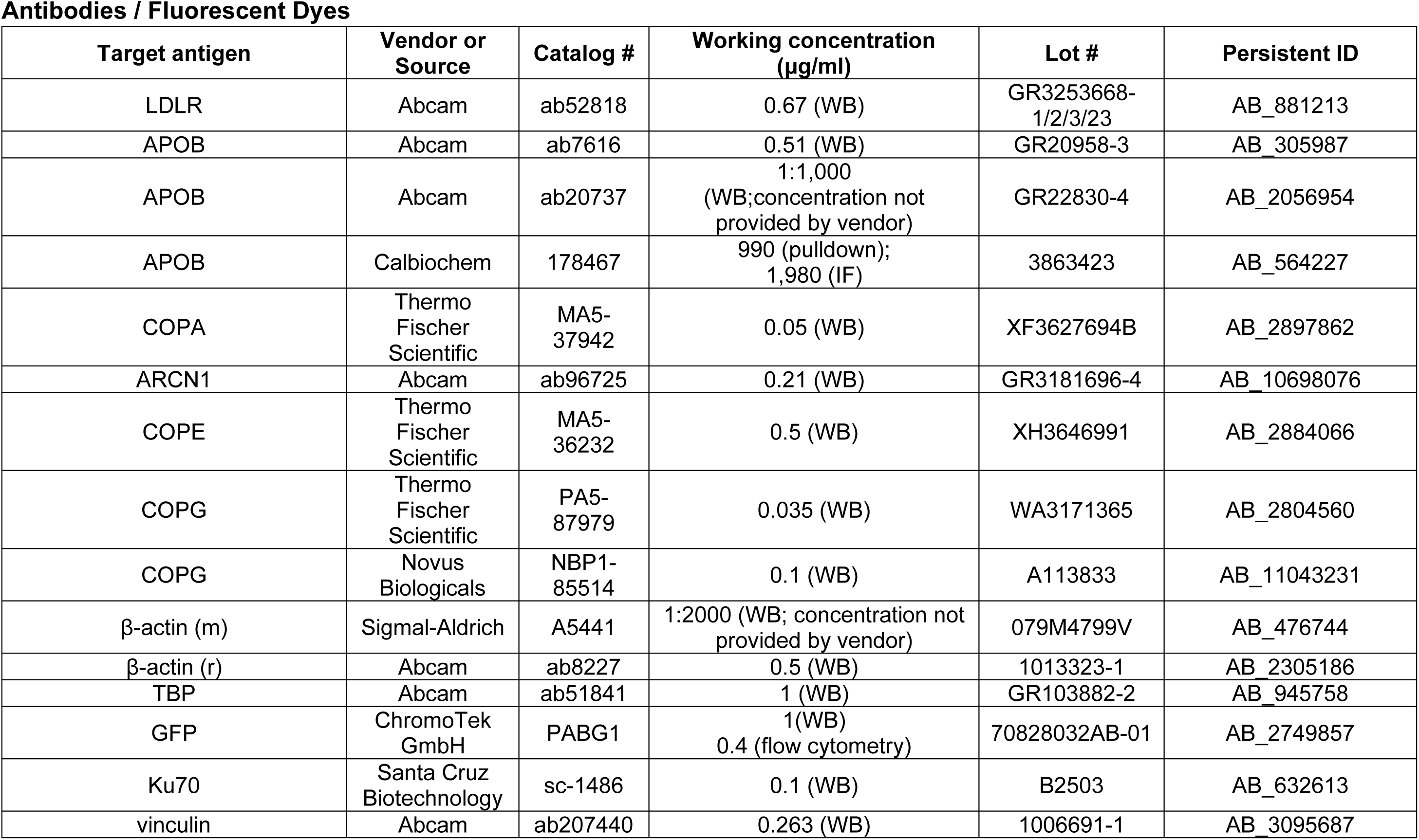
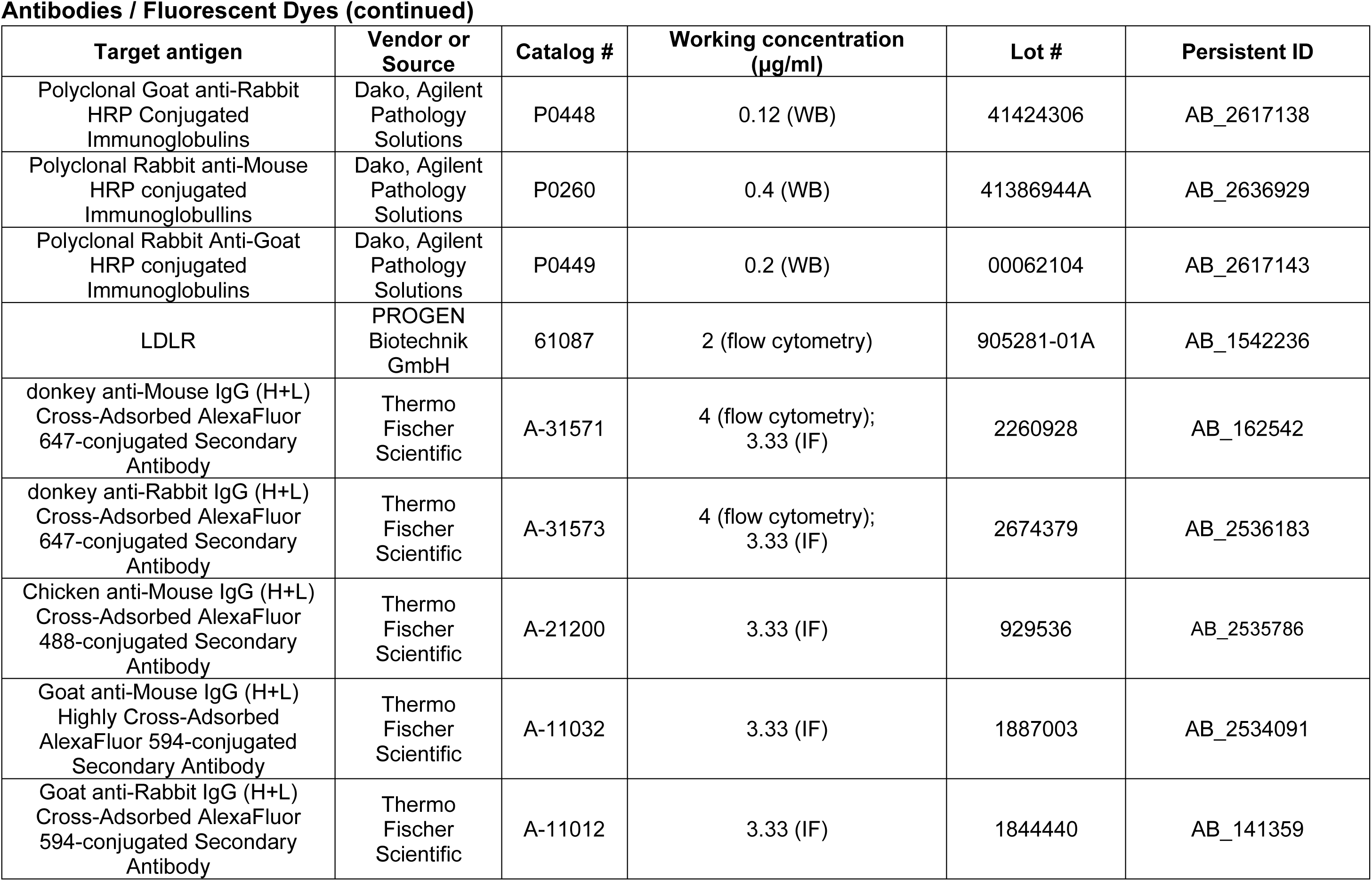
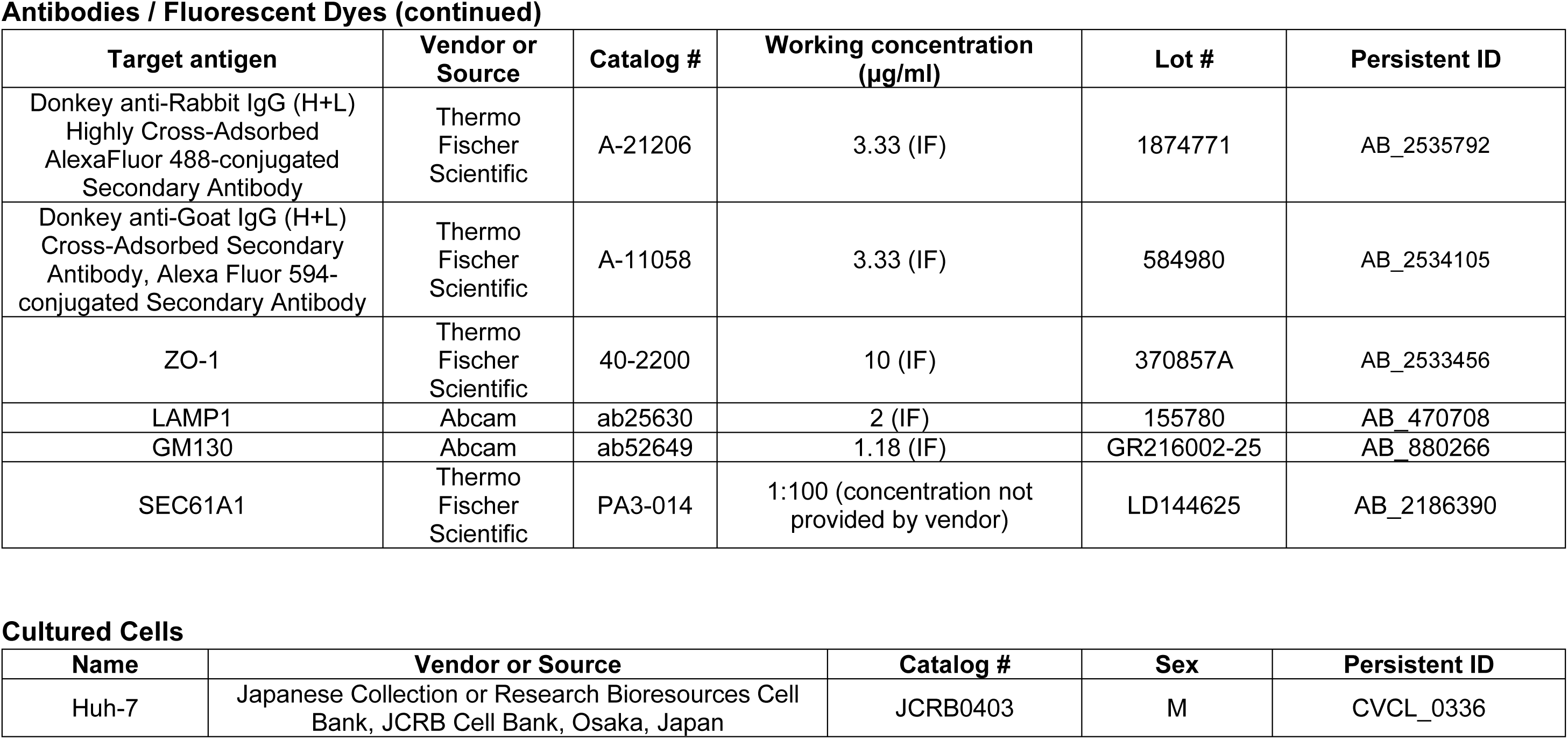
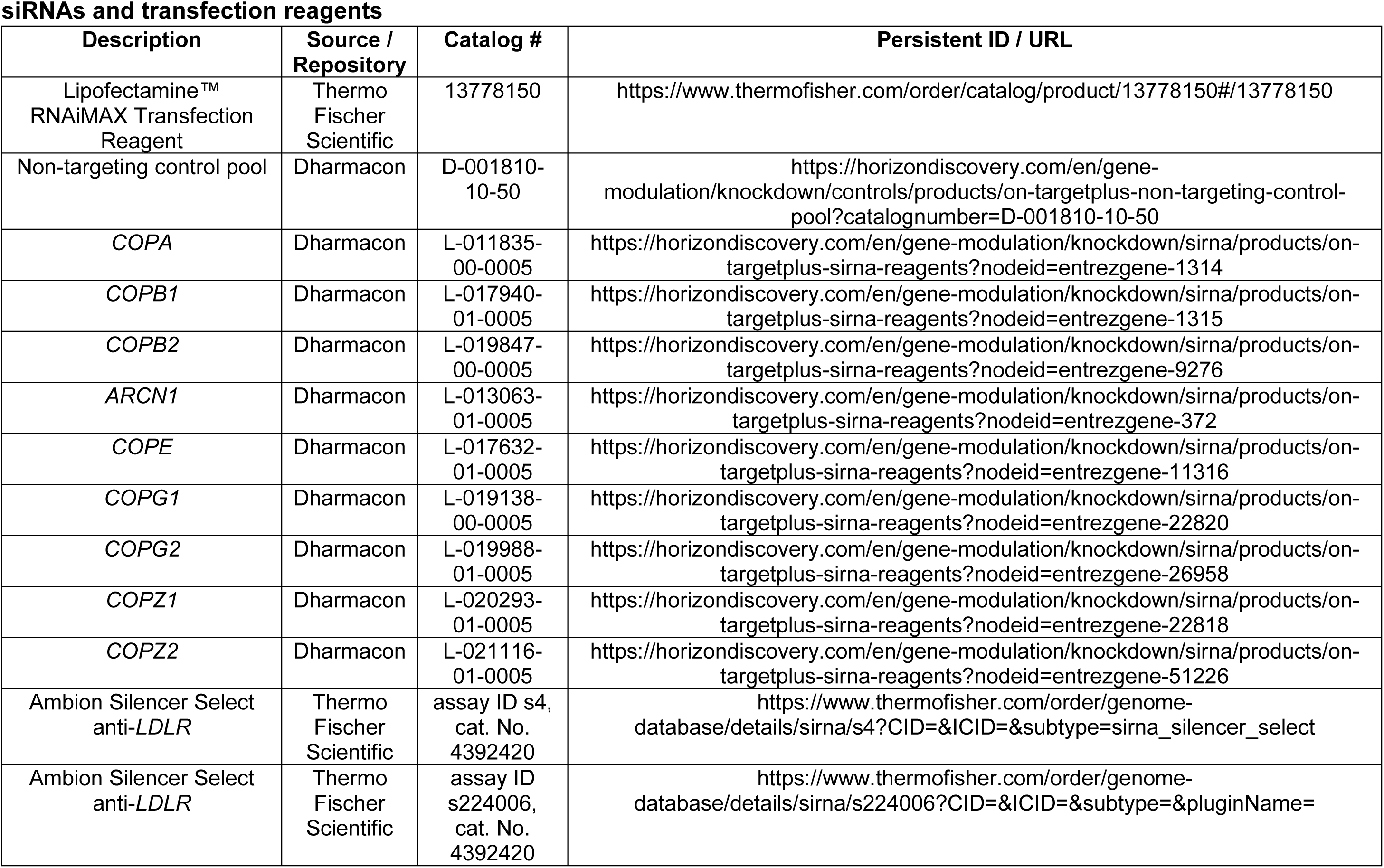
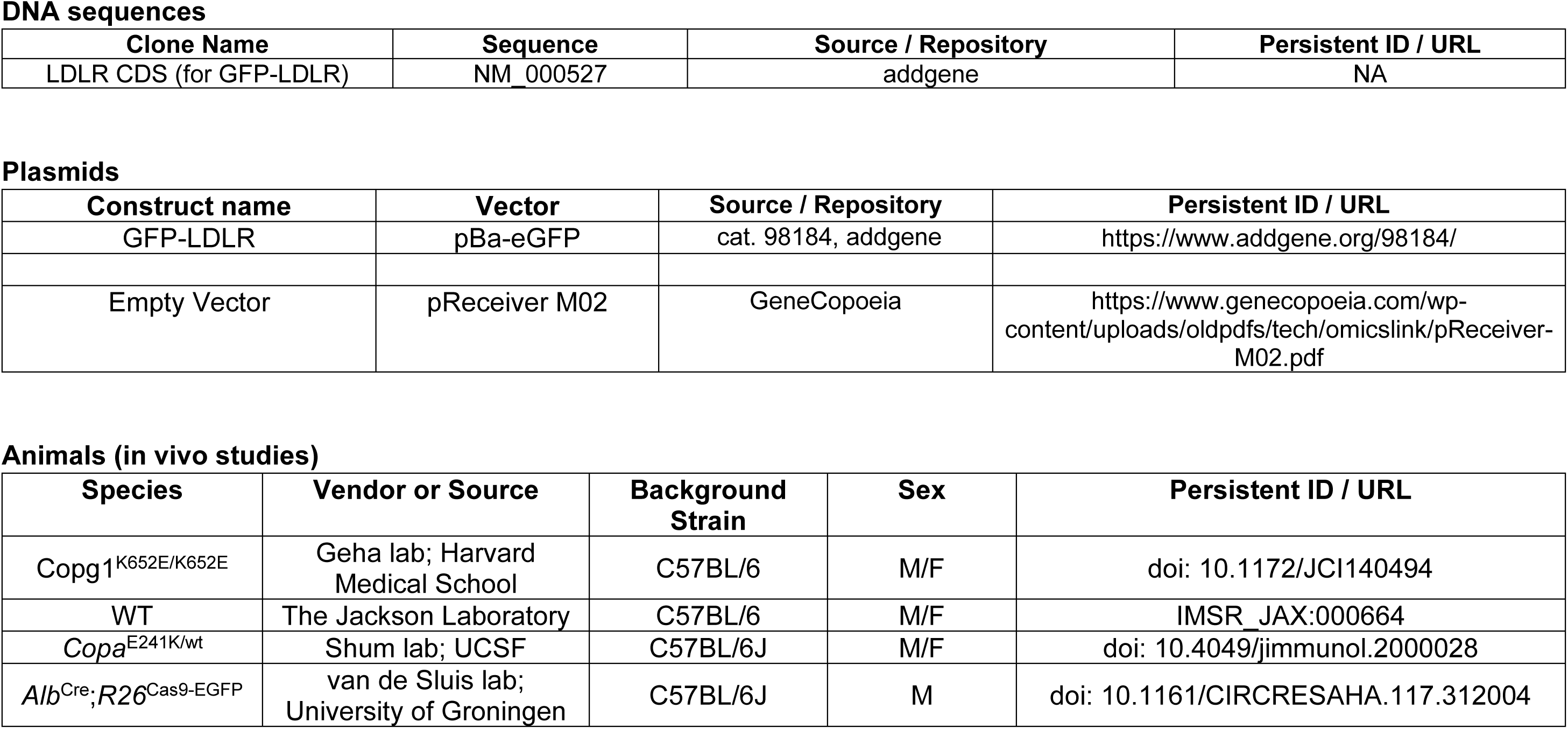
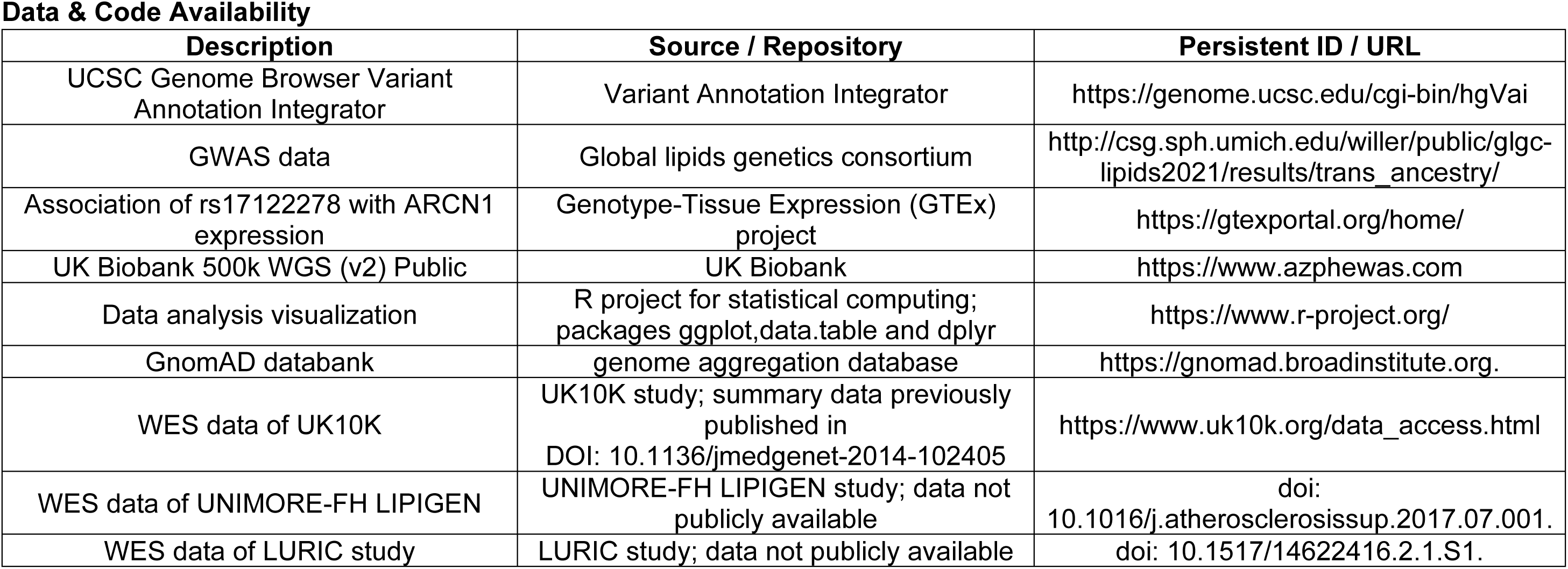

Antibodies / Fluorescent Dyes
| Target antigen | Vendor or Source | Catalog # | Working concentration (µg/ml) | Lot # | Persistent ID |
| --- | --- | --- | --- | --- | --- |
| LDLR | Abcam | ab52818 | 0.67 (WB) | GR3253668-1/2/3/23 | AB_881213 |
| APOB | Abcam | ab7616 | 0.51 (WB) | GR20958-3 | AB_305987 |
| APOB | Abcam | ab20737 | 1:1,000<br>(WB;concentration not provided by vendor) | GR22830-4 | AB_2056954 |
| APOB | Calbiochem | 178467 | 990 (pulldown);<br>1,980 (IF) | 3863423 | AB_564227 |
| COPA | Thermo Fischer Scientific | MA5-37942 | 0.05 (WB) | XF3627694B | AB_2897862 |
| ARCN1 | Abcam | ab96725 | 0.21 (WB) | GR3181696-4 | AB_10698076 |
| COPE | Thermo Fischer Scientific | MA5-36232 | 0.5 (WB) | XH3646991 | AB_2884066 |
| COPG | Thermo Fischer Scientific | PA5-87979 | 0.035 (WB) | WA3171365 | AB_2804560 |
| COPG | Novus Biologicals | NBP1-85514 | 0.1 (WB) | A113833 | AB_11043231 |
| β-actin (m) | Sigma-Aldrich | A5441 | 1:2000 (WB; concentration not provided by vendor) | 079M4799V | AB_476744 |
| β-actin (r) | Abcam | ab8227 | 0.5 (WB) | 1013323-1 | AB_2305186 |
| TBP | Abcam | ab51841 | 1 (WB) | GR103882-2 | AB_945758 |
| GFP | ChromoTek GmbH | PABG1 | 1(WB)<br>0.4 (flow cytometry) | 70828032AB-01 | AB_2749857 |
| Ku70 | Santa Cruz Biotechnology | sc-1486 | 0.1 (WB) | B2503 | AB_632613 |
| vinculin | Abcam | ab207440 | 0.263 (WB) | 1006691-1 | AB_3095687 |
| Polyclonal Goat anti-Rabbit HRP Conjugated Immunoglobulins | Dako, Agilent Pathology Solutions | P0448 | 0.12 (WB) | 41424306 | AB_2617138 |
| Polyclonal Rabbit anti-Mouse HRP conjugated Immunoglobulins | Dako, Agilent Pathology Solutions | P0260 | 0.4 (WB) | 41386944A | AB_2636929 |
| Polyclonal Rabbit Anti-Goat HRP conjugated Immunoglobulins | Dako, Agilent Pathology Solutions | P0449 | 0.2 (WB) | 00062104 | AB_2617143 |
| LDLR | PROGEN Biotechnik GmbH | 61087 | 2 (flow cytometry) | 905281-01A | AB_1542236 |
| donkey anti-Mouse IgG (H+L) Cross-Adsorbed AlexaFluor 647-conjugated Secondary Antibody | Thermo Fischer Scientific | A-31571 | 4 (flow cytometry); 3.33 (IF) | 2260928 | AB_162542 |
| donkey anti-Rabbit IgG (H+L) Cross-Adsorbed AlexaFluor 647-conjugated Secondary Antibody | Thermo Fischer Scientific | A-31573 | 4 (flow cytometry); 3.33 (IF) | 2674379 | AB_2536183 |
| Chicken anti-Mouse IgG (H+L) Cross-Adsorbed AlexaFluor 488-conjugated Secondary Antibody | Thermo Fischer Scientific | A-21200 | 3.33 (IF) | 929536 | AB_2535786 |
| Goat anti-Mouse IgG (H+L) Highly Cross-Adsorbed AlexaFluor 594-conjugated Secondary Antibody | Thermo Fischer Scientific | A-11032 | 3.33 (IF) | 1887003 | AB_2534091 |
| Goat anti-Rabbit IgG (H+L) Cross-Adsorbed AlexaFluor 594-conjugated Secondary Antibody | Thermo Fischer Scientific | A-11012 | 3.33 (IF) | 1844440 | AB_141359 |
| Donkey anti-Rabbit IgG (H+L) Highly Cross-Adsorbed AlexaFluor 488-conjugated Secondary Antibody | Thermo Fischer Scientific | A-21206 | 3.33 (IF) | 1874771 | AB_2535792 |
| Donkey anti-Goat IgG (H+L) Cross-Adsorbed Secondary Antibody, Alexa Fluor 594-conjugated Secondary Antibody | Thermo Fischer Scientific | A-11058 | 3.33 (IF) | 584980 | AB_2534105 |
| ZO-1 | Thermo Fischer Scientific | 40-2200 | 10 (IF) | 370857A | AB_2533456 |
| LAMP1 | Abcam | ab25630 | 2 (IF) | 155780 | AB_470708 |
| GM130 | Abcam | ab52649 | 1.18 (IF) | GR216002-25 | AB_880266 |
| SEC61A1 | Thermo Fischer Scientific | PA3-014 | 1:100 (concentration not provided by vendor) | LD144625 | AB_2186390 |

**Cultured Cells**
| Name | Vendor or Source | Catalog # | Sex | Persistent ID |
| --- | --- | --- | --- | --- |
| Huh-7 | Japanese Collection or Research Bioresources Cell Bank, JCRB Cell Bank, Osaka, Japan | JCRB0403 | M | CVCL_0336 |

siRNAs and transfection reagents
| Description | Source / Repository | Catalog # | Persistent ID / URL |
| --- | --- | --- | --- |
| Lipofectamine™ RNAiMAX Transfection Reagent | Thermo Fischer Scientific | 13778150 | https://www.thermofisher.com/order/catalog/product/13778150#/13778150 |
| Non-targeting control pool | Dharmacon | D-001810-10-50 | https://horizondiscovery.com/en/gene-modulation/knockdown/controls/products/on-targetplus-non-targeting-control-pool?catalognumber=D-001810-10-50 |
| *COPA* | Dharmacon | L-011835-00-0005 | https://horizondiscovery.com/en/gene-modulation/knockdown/sirna/products/on-targetplus-sirna-reagents?nodeid=entrezgene-1314 |
| *COPB1* | Dharmacon | L-017940-01-0005 | https://horizondiscovery.com/en/gene-modulation/knockdown/sirna/products/on-targetplus-sirna-reagents?nodeid=entrezgene-1315 |
| *COPB2* | Dharmacon | L-019847-00-0005 | https://horizondiscovery.com/en/gene-modulation/knockdown/sirna/products/on-targetplus-sirna-reagents?nodeid=entrezgene-9276 |
| *ARCN1* | Dharmacon | L-013063-01-0005 | https://horizondiscovery.com/en/gene-modulation/knockdown/sirna/products/on-targetplus-sirna-reagents?nodeid=entrezgene-372 |
| *COPE* | Dharmacon | L-017632-01-0005 | https://horizondiscovery.com/en/gene-modulation/knockdown/sirna/products/on-targetplus-sirna-reagents?nodeid=entrezgene-11316 |
| *COPG1* | Dharmacon | L-019138-00-0005 | https://horizondiscovery.com/en/gene-modulation/knockdown/sirna/products/on-targetplus-sirna-reagents?nodeid=entrezgene-22820 |
| *COPG2* | Dharmacon | L-019988-01-0005 | https://horizondiscovery.com/en/gene-modulation/knockdown/sirna/products/on-targetplus-sirna-reagents?nodeid=entrezgene-26958 |
| *COPZ1* | Dharmacon | L-020293-01-0005 | https://horizondiscovery.com/en/gene-modulation/knockdown/sirna/products/on-targetplus-sirna-reagents?nodeid=entrezgene-22818 |
| *COPZ2* | Dharmacon | L-021116-01-0005 | https://horizondiscovery.com/en/gene-modulation/knockdown/sirna/products/on-targetplus-sirna-reagents?nodeid=entrezgene-51226 |
| Ambion Silencer Select anti-*LDLR* | Thermo Fischer Scientific | assay ID s4, cat. No. 4392420 | https://www.thermofisher.com/order/genome-database/details/sirna/s4?CID=&ICID=&subtype=sirna_silencer_select |
| Ambion Silencer Select anti-*LDLR* | Thermo Fischer Scientific | assay ID s224006, cat. No. 4392420 | https://www.thermofisher.com/order/genome-database/details/sirna/s224006?CID=&ICID=&subtype=&pluginName= |

DNA sequences
| Clone Name | Sequence | Source / Repository | Persistent ID / URL |
| --- | --- | --- | --- |
| LDLR CDS (for GFP-LDLR) | NM_000527 | addgene | NA |

Plasmids
| Construct name | Vector | Source / Repository | Persistent ID / URL |
| --- | --- | --- | --- |
| GFP-LDLR | pBa-eGFP | cat. 98184, addgene | https://www.addgene.org/98184/ |
| Empty Vector | pReceiver M02 | GeneCopoeia | https://www.genecopoeia.com/wp-content/uploads/oldpdfs/tech/omicslink/pReceiver-M02.pdf |

Animals (in vivo studies)
| Species | Vendor or Source | Background Strain | Sex | Persistent ID / URL |
| --- | --- | --- | --- | --- |
| Copg1<sup>K652E/K652E</sup> | Geha lab; Harvard Medical School | C57BL/6 | M/F | doi: 10.1172/JCI140494 |
| WT | The Jackson Laboratory | C57BL/6 | M/F | IMSR_JAX:000664 |
| Copa<sup>E241K/wt</sup> | Shum lab; UCSF | C57BL/6J | M/F | doi: 10.4049/jimmunol.2000028 |
| Alb<sup>Cre</sup>;R26<sup>Cas9-EGFP</sup> | van de Sluis lab; University of Groningen | C57BL/6J | M | doi: 10.1161/CIRCRESAHA.117.312004 |

**Data & Code Availability**
| **Description** | **Source / Repository** | **Persistent ID / URL** |
| --- | --- | --- |
| UCSC Genome Browser Variant Annotation Integrator | Variant Annotation Integrator | https://genome.ucsc.edu/cgi-bin/hgVai |
| GWAS data | Global lipids genetics consortium | http://csg.sph.umich.edu/willer/public/glgc-lipids2021/results/trans_ancestry/ |
| Association of rs17122278 with ARCN1 expression | Genotype-Tissue Expression (GTEx) project | https://gtexportal.org/home/ |
| UK Biobank 500k WGS (v2) Public | UK Biobank | https://www.azphewas.com |
| Data analysis visualization | R project for statistical computing; packages ggplot, data.table and dplyr | https://www.r-project.org/ |
| GnomAD databank | genome aggregation database | https://gnomad.broadinstitute.org. |
| WES data of UK10K | UK10K study; summary data previously published in<br>DOI: 10.1136/jmedgenet-2014-102405 | https://www.uk10k.org/data_access.html |
| WES data of UNIMORE-FH LIPIGEN | UNIMORE-FH LIPIGEN study; data not publicly available | doi:<br>10.1016/j.atherosclerosissup.2017.07.001. |
| WES data of LURIC study | LURIC study; data not publicly available | doi: 10.1517/14622416.2.1.S1. |

